# Multiplexed Glycan Immunofluorescence Identification of Pancreatic Cancer Cell Subpopulations in Both Tumor and Blood Samples

**DOI:** 10.1101/2024.08.22.609143

**Authors:** Braelyn Binkowski, Zachary Klamer, ChongFeng Gao, Ben Staal, Anna Repesh, Hoang-Le Tran, David M. Brass, Pamela Bartlett, Steven Gallinger, Maria Blomqvist, J. Bradley Morrow, Peter Allen, Chanjuan Shi, Aatur Singhi, Randall Brand, Ying Huang, Galen Hostetter, Brian B. Haab

## Abstract

Pancreatic ductal adenocarcinoma (PDAC) tumor heterogeneity impedes the development of biomarker assays suitable for early disease detection that would improve patient outcomes. The CA19-9 glycan is currently used as a standalone biomarker for PDAC. Furthermore, previous studies have shown that cancer cells may display aberrant membrane-associated glycans. We therefore hypothesized that PDAC cancer cell subpopulations could be distinguished by aberrant glycan signatures. We used multiplexed glycan immunofluorescence combined with pathologist annotation and automated image processing to distinguish between PDAC cancer cell subpopulations within tumor tissue. Using a training-set/test-set approach, we found that PDAC cancer cells may be identified by signatures comprising 4 aberrant glycans (VVL, CA19-9, sTRA, and GM2) and that there are three glycan-defined PDAC tumor types: sTRA type, CA19-9 type, and intermixed. To determine whether the aberrant glycan signatures could be detected in blood samples, we developed hybrid glycan sandwich assays for membrane-associated glycans. In both patient-matched tumor and blood samples, the proportion of aberrant glycans detected was consistent. Furthermore, our multiplexed glycan immunofluorescent approach proved to be more sensitive and more specific than CA19-9 alone. Our results provide proof of concept for a novel methodology to improve early PDAC detection and patient outcomes.

## Introduction

The early detection and treatment of pancreatic ductal adenocarcinoma (PDAC) results in better patient outcomes for this extremely aggressive cancer, but only ∼15% of patients are diagnosed at a resectable stage (IA, IB, or IIA) where the tumor has not yet spread to major blood vessels, regional lymph nodes, or distant organs (1). There are currently no biomarker assays that will identify PDAC at an early, localized stage when curative resection is an option. A barrier to the development of such assays is the heterogeneity between PDAC tumors in the subpopulations of cancer cells they contain. Previous studies revealed that PDAC tumors diverge widely in the intratumor makeup of the PDAC subpopulations and macromolecular compositions. Two primary PDAC cancer cell subpopulations, termed classical and basal-like (or squamous), have been identified by gene expression profiling on whole-tumor mRNA (2–4), and other, less-well-characterized PDAC subpopulations have been identified by single-cell and spatial transcriptomics analyses (5, 6). One study divided the primary subtypes into five categories (5), while another categorized the cancer cells by their subclonal lineages and their states (cycling, adhesive, *etc*.) (6). Some tumors contain mostly basal-like cancer cells or mostly classical cancer cells, while other tumors have a mix of the two (7, 8), together with varying numbers of cancer cells that have both classical and basal-like characteristics (7, 9, 10). Because of this heterogeneity, no single biomarker assay identifies PDACs across their range of subpopulation makeup.

A potential solution to this problem is to use assays that are more specific to each subpopulation in combination to complementarily detect the individual cancer cells subpopulations that make up PDAC tumors. Various approaches have been developed for detecting different cancer cell subpopulations within PDAC tumors. These assays include a 16 gene signature (11), multi-protein immunochemical assays (7, 12, 13), and individual immunochemical assays used to detect GATA6 (14) and KRT17 (15, 16). Such assays could be useful in clinical applications to differentiate between cancer cell subpopulations and guide treatment, such as to predict differential drug responses between the classical and basal-like subtypes (11, 12). The assays, however, are not suitable for the diagnostic use of subpopulation-linked biomarkers for the complementary detection of divergent tumors, as they are not readily detectable in an easily accessible specimen such as blood plasma. An additional challenge is the potential detection of non-cancer cells and resulting false-positive identifications of the current protein biomarkers. The previous studies did not determine detection sensitivity relative to all cancer cells or the detection specificity relative to non-cancer cells, although multiple types of non-cancer cells can also express the transcription factor and keratin protein biomarkers that have been identified. Therefore, we used an approach based on molecular patterns that could distinguish between PDAC subpopulations and non-cancer cells and that could be detected in blood samples (serum or plasma).

All cells are decorated with complex carbohydrate structures known as glycans. The cell surface glycome, or the complete array of glycans on cell membranes, results through complex biosynthetic processes involving multiple enzymes, substrates, and metabolites. As such, the cell surface glycome is not linked to the expression levels of any gene or protein but is instead regulated by multiple inputs that are specific to each cell type. This specificity is illustrated by glycans that are already widely used as cell type markers, such as CD15 for neutrophils, CD65s for myeloid differentiation, and TRA-1-60 for induced pluripotent stem cells, among others. As cells transform from normal cells to abnormal dysplastic cells and then to cancer cells, they activate biological programs of gene expression and metabolism that are not present in the originating cells and that affect the biosynthesis of glycans (13–15). PDAC cells have glycosylation distinct from non-cancer cells and between subsets of PDAC cells, as shown by MALDI glycan imaging (16). Furthermore, cancer-associated glycans can be detected using immunochemical assays on both cancer cell surfaces and on cellular secretions in the peripheral blood plasma (17), making them ideal for improved cancer identification in surveillance and monitoring applications. Therefore, identifying how the cancer glycome is composed and develops could increase our ability to identify and stratify early stage PDAC patients by their unique glycan signatures.

CA19-9 and sTRA are established PDAC biomarkers that provided impetus to define glycan biomarkers of PDAC subpopulations. CA19-9 is used as a plasma biomarker to confirm diagnoses and monitor the trajectory of PDAC (18), but at least half of PDAC cancer cell subpopulations in tumors do not produce CA19-9 (19). The sTRA glycan is structurally related to CA19-9 but produced by different subpopulations of PDAC cells, so that separate subpopulations of PDAC cells display one, both, or neither of these two glycans. The CA19-9 exclusive cells are epithelial and mostly classical, while the sTRA exclusive cells include both epithelial/classical and mesenchymal/basal-like (20), indicating a general but not exact correspondence between expression-based and glycan-based classifications. Owing to the complementarity of the glycans, a subset of tumors may be detected with CA19-9 while another subset is better detected with sTRA. Indeed, the plasma detection of CA19-9 in combination with sTRA identified significantly more PDAC patients than by either individual biomarker alone, while maintaining a matched low rate of false-positive identification (17, 20–23). Therefore, the cancer glycome and the signatures of subpopulation-specific glycans may better identify diverse PDAC tumors than any other approach.

We explored whether distinct PDAC subpopulations display different cancer-associated glycan signatures that enable their specific detection. We approached this question using multiplexed glycan immunofluorescence on primary human PDAC tumors. To our knowledge, this study is the first analysis of cancer-associated glycan signatures on specific cell populations in primary tumors. To enable the identification of glycan signatures that could identify PDAC cell subpopulations, we developed a pipeline consisting of multiplexed glycan immunofluorescence, automated thresholding, and an analysis method that connects glycan profiles with cell types. We determined that PDAC cell subpopulations display aberrant signatures of glycans that rarely occur on non-cancer cells. Tumors display distinct types of intratumoral heterogeneity in their glycan-defined PDAC subpopulations. Furthermore, these glycan signatures could be detected in the peripheral blood, improving the sensitivity and specificity of noninvasive PDAC identification through detecting tumors with heterogeneous subpopulations of cancer cells.

## Results

### PDAC Cells Display Aberrant Glycan Signatures

To begin to address the challenge presented by the heterogeneity of PDAC cancer cell subpopulations, we asked whether the glycans that decorate the cellular plasma membrane— the glycan signature of the cell—differ between PDAC cancer cells and non-cancer cells (**Fig. 1A**). To develop optimized assays for each of these glycan motifs, we analyzed glycan microarray data from the CarboGrove database (24) to select glycan-binding antibodies and lectins (glycan-binding proteins, GBPs) (**Fig. 1B, Supplementary Table 1**) that specifically bind to PDAC-associated glycan motifs in addition to CA19-9 and sTRA. For example, the Tn antigen is overproduced in many cancers including PDAC (25, 26), glycan motifs with a terminal GlcNAc—as detected with monoclonal antibodies or GBPs such as GSL-II (16, 27)—have previously been linked to subsets of PDAC cells, the TRA-1-60 epitope was found as a marker of chemoresistant, stem-like PDAC cells (28, 29), and the GM2 ganglioside—a glycolipid typically associated with neuroendocrine cells—has been identified on PDAC cells with neuroendocrine characteristics (30). These glycans have been investigated individually in PDAC tissues but never in combination. We therefore asked whether individual cancer-associated glycan motifs considered together could better distinguish cancer from non-cancer than any single biomarker alone.

**Figure 1.**
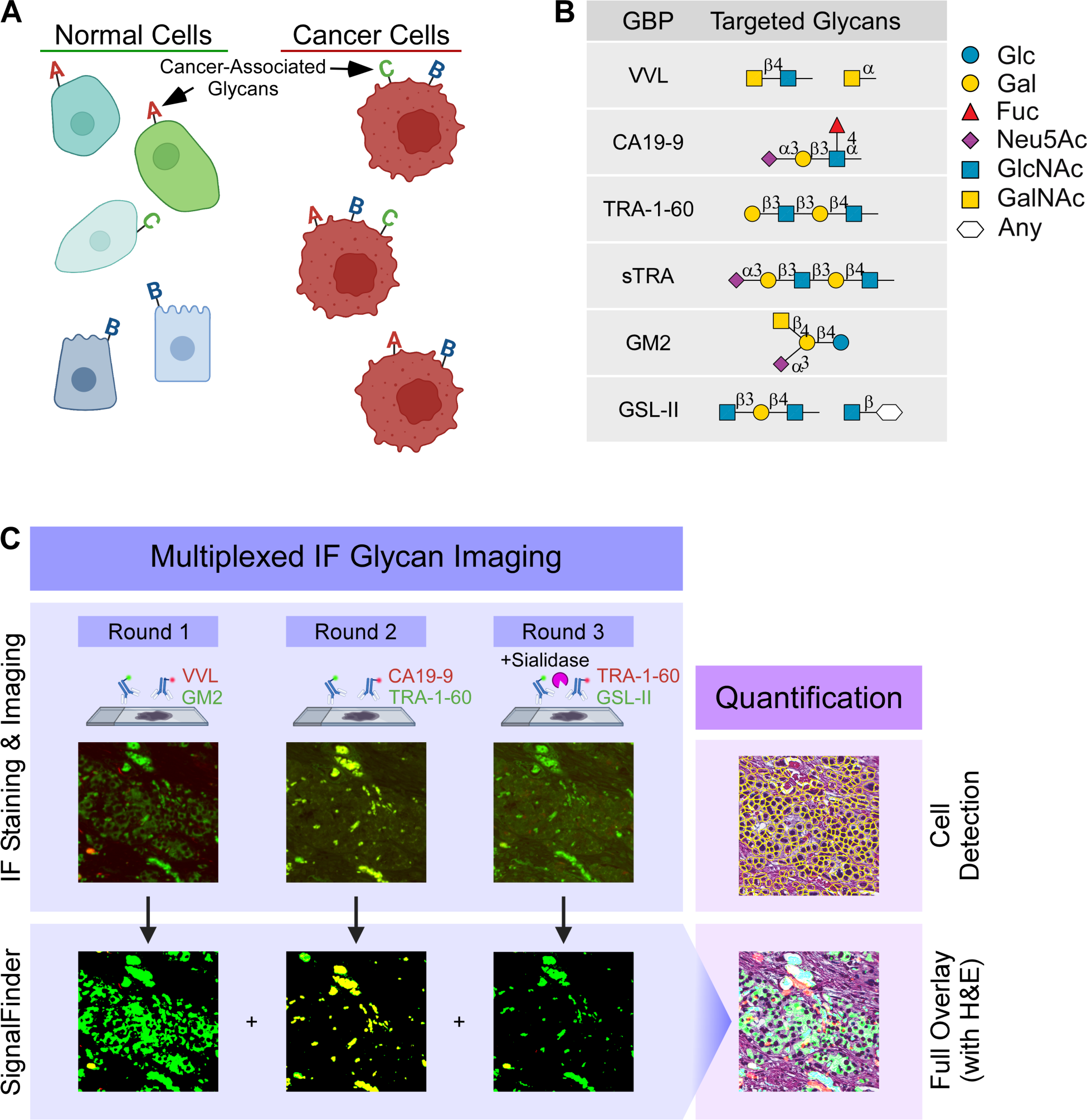
Identification and quantification of cellular glycan signatures. A) Model of cell surface glycan signatures. B) Glycan-binding proteins (GBP) and their targets. C) Multiplexed immunofluorescence with automated signal thresholding by SignalFinder, followed by assignments of glycan signatures to individual cells.

We then used the GBPs that were selected using the CarboGrove data to localize their target glycans in whole block primary tissue specimens using multiplexed glycan immunofluorescence as shown in **Figure 1C**. We used SignalFinder to automate the thresholding of primary image data (16, 28, 29). Automated thresholding enables intervention-free capture of immunofluorescent images from tumor tissue samples that are suitable for objective, consistent, and unbiased comparisons of signal intensity between tumor specimens. The output of SignalFinder—a binary map of pixels that are above the background (rather than a gradient of signal)—was aligned with the brightfield image of the H&E-stained tissue using the Warpy FiJI plugin (31, 32) **Fig. 1C**). We then quantified the percent of each cell within each region of interest (ROI) that was positive for each glycan.

We analyzed whole block sections from 34 PDAC tumors and 4 non-cancerous pancreas specimens (**Table 1 and Supplementary Table 2**). We used whole block sections to enable comparisons across the wide range of PDAC and non-cancer cell subpopulations in PDAC tumors. In a training set of 15 PDAC tumors, we tested the associations between the glycan signatures and cell types annotated by the study pathologist (G.H.) (**Fig. 2A and Supplementary Figure 1**). The pathologist annotated 13 types of histological features (**Supplementary Table 3 and Supplementary Figure 1)** within ROIs on the brightfield images of the H&E-stained tissue sections, which contained 39 to 111 ROIs in each tumor for a total of 1149 ROIs and 666,544 detected cells. We asked whether we could distinguish between cells in the cancer ROIs and cells in the non-cancer ROIs based on the presence of any single glycan alone or the presence of an aberrant permutation, or signature, of glycans. We used recursive partitioning to ask whether specific subpopulations of cancer cells preferentially display different glycan signatures more frequently than the non-cancer cells. The analysis showed that 4 detections—VVL, CA19-9, sTRA, and GM2— were associated with the cancer cells (**Supplementary Figure 2)**. To enable delineation of the associations between specific glycan signatures and cancer cells, we specified the glycan signature of each cell by classifying each cell as positive or negative for each of the four cancer-associated glycans and then using a four-digit binary code to represent the glycan signature. To classify the positivity or negativity of each glycan in each cell, we used the cutoffs in the fraction-positive values for each glycan used in the classification tree (**Supplementary Figure 2**). In this binary code system, the glycans retain the order VVL, CA19-9, sTRA, and GM2. Therefore, the code 0100 indicates that CA19-9 alone is displayed by a cell. We used these analyses to quantify the percentages of cells in each ROI that are positive for each glycan signature as represented by the four-digit binary code (**Fig. 2B**).

**Figure 2.**
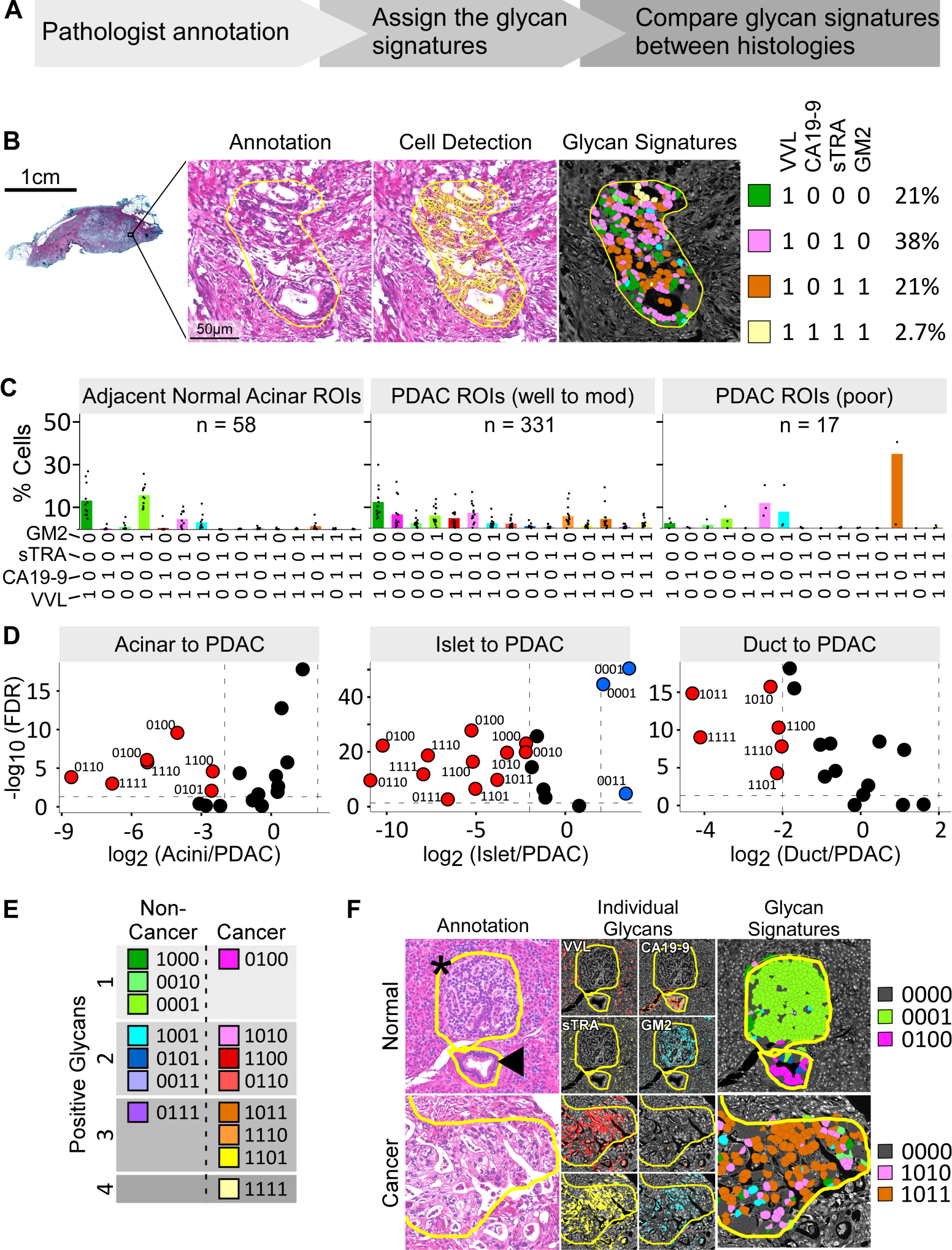
Training set identification of cancer-association glycan signatures. A) Training design. B) Quantification of cells with each glycan signature in ROIs. C) Distributions of cell surface glycan signatures among acinar and PDAC cells. D) Volcano plots of pairwise comparisons between cell types. E) Cancer-associated and non-cancer-associated glycan signatures. F) Representative images and quantifications of non-cancer and cancer ROIs. The asterisk marks normal islet cells and arrowhead marks a normal duct.

**Table 1.** Tumor specimens. N/A, not available; -, not applicable; *, 2 specimens obtained from 1 patient.

|  |  | Duke | UHN | UPMC | VAI | Total |
| --- | --- | --- | --- | --- | --- | --- |
| Specimens |  | 21 | 4 | 9 | 4 | 38 |
| Sex | Male | 9 | 2 | 5 | 2* | 16 |
|  | Female | 12 | 2 | 4 | 1 | 18 |
| Age at Dx | Mean | 66.48 | 57.75 | 71 | - | 66.65 |
|  | SD | 10.66 | 15.39 | 8.34 | - | 11.05 |
|  | Median | 68 | 55.5 | 73 | - | 68.5 |
|  | Min | 39 | 43 | 56 | - | 39 |
|  | Max | 80 | 77 | 83 | - | 83 |
| Specimen Type | PDAC | 21 | 4 | 9 | 0 | 34 |
|  | Normal | 0 | 0 | 0 | 4 | 4 |
| Site | Head | 14 | N/A | 7 | - |  |
|  | Tail | 7 | N/A | 2 | - |  |
| Margins | Positive | 7 | 0 | 2 | - | 9 |
|  | Negative | 15 | 4 | 7 | - | 26 |
| Lymph Nodes | Positive | 13 | 1 | 2 | - | 16 |
|  | Negative | 8 | 3 | 7 | - | 18 |
| Neoadjuvant Treated | Yes | 0 | 0 | 9 | - | 9 |
|  | No | 21 | 4 | 0 | - | 29 |

Using our four-digit binary code, we analyzed the signatures of the glycans displayed on the cells in the analyzed ROIs. We found that cells in the non-cancer ROIs were predominantly positive for a single glycan alone. For example, cells in acinar ROIs were predominantly VVL alone (code 1000) or GM2 alone (code 0001) (**Fig. 2C**). Islet cells were predominantly GM2 alone (code 0001), and ductal cells were predominantly VVL, CA19-9, or GM2 alone (codes 1000, 0100, and 0001, respectively) **Supplementary Figure 3**). These signatures were similar between ROIs adjacent to the tumors and ROIs in healthy pancreas samples (**Supplementary Figure 3**). In contrast to the non-cancer ROIs, the PDAC ROIs included a broader mix of glycan signatures, including cells that produced signatures of 2, 3, or 4 glycans (**Fig. 2C**). To identify glycan signatures that associate with the cancer cells, we used a classification tree to compare the dichotomized glycan values of the cells in the cancer ROIs to the cells in the non-cancer ROIs. We found that eight different glycan signatures were more frequently on cancer cells that on non-cancer cells (**Supplementary Figure 2**). Seven were positive for 2-4 glycans and one was positive for only 1 glycan (CA19-9, code 0100) (**Supplementary Figure 2**).

In direct comparisons of the cancer to the non-cancer ROIs, all 8 of the signatures identified above were significantly associated with PDAC (p < 0.05, Wilcoxon test, FDR correction) (**Fig. 2D, 2E**). All 8 of the glycan signatures also had a significantly higher probability (p < 0.05, log odds of probability) of belonging to a cancer ROI than to a normal ductal, acinar, or islet ROI (**Supplementary Figure 3).** Five of the non-cancer associated glycan signatures (**Supplementary Figure 2**) were higher in the PDAC ROIs when compared to non-cancer ROIs (code 0101 in the acinar to PDAC comparison and codes 1000, 0010, 0101, and 0111 in the islet to PDAC comparison, **Fig. 2D and Supplementary Figure 3**). However, these 5 glycan signatures varied between the corresponding non-cancer cell types (**Supplementary Figure 3**) and were ultimately considered indicative of specific normal cells rather than of PDAC cells. Representative images (**Fig. 2F** and **Supplementary Figure 4**) indicated that ductal cells were mainly positive for CA19-9 alone and islet cells were positive mainly for GM2 alone, and that the cancer cells were positive for 2 or 3 of the glycans.

To determine whether we could use these 2-4 glycan signatures to differentiate between cancer and non-cancer in PDAC tissues, we analyzed a test set of the remaining 19 PDAC specimens and 4 non-cancer specimens (**Fig. 3A**). We first identified regions in each tumor with spatially dense clusters of each of the glycan signatures based on the method described earlier (21). This method calculates the spatial density of each glycan signal and converts the overlap of the spatially dense regions to distinct ROIs with the corresponding four-digit binary code. We randomly sampled the ROIs at an average of 112 ROIs per tissue (range 20 -165), for a total of 2,135 ROIs (average 308 cells/ROI) (**Supplementary Table 4**). Using annotations made on the images of the H&E-stained tumors by a pathologist (G.H.) who was blinded to the glycan data, we compared the prevalence of each glycan signature between the annotated cancer ROIs (combined well, moderate, and poor differentiation, 461 ROIs) and the non-cancer ROIs (lymphoid, islet, acini, connective tissue, peripheral nerve, blood vessels, normal duct, macrophages, 791 ROIs) (**Supplementary Table 4 and complete list in Supplementary Table 5**). We excluded ROIs with tissue-fold artifacts, high levels of debris, or non-pancreatic features. Each cancer-associated glycan signature was represented in ROIs from multiple tumors, with variation between the tumors in types and variety of glycan signatures (**Fig. 3B**). By classifying each ROI by CA19-9 alone as positive or negative using the same cutoff as used in the multi-glycan analysis, we observed that some of the CA19-9-negative cancer ROIs were positive for other cancer-associated glycan signatures (as in tumors 10-16673 and 18-137) (**Fig. 3B**). Three of the 19 tumors were completely negative for CA19-9 (18-460, 33524, and 09-5741); all three had cancer ROIs that were positive for putative cancer-associated glycan signatures (41-87% of the cancer ROIs) with 0% positivity in the non-cancer ROIs (**Fig. 3B**).

**Figure 3.**
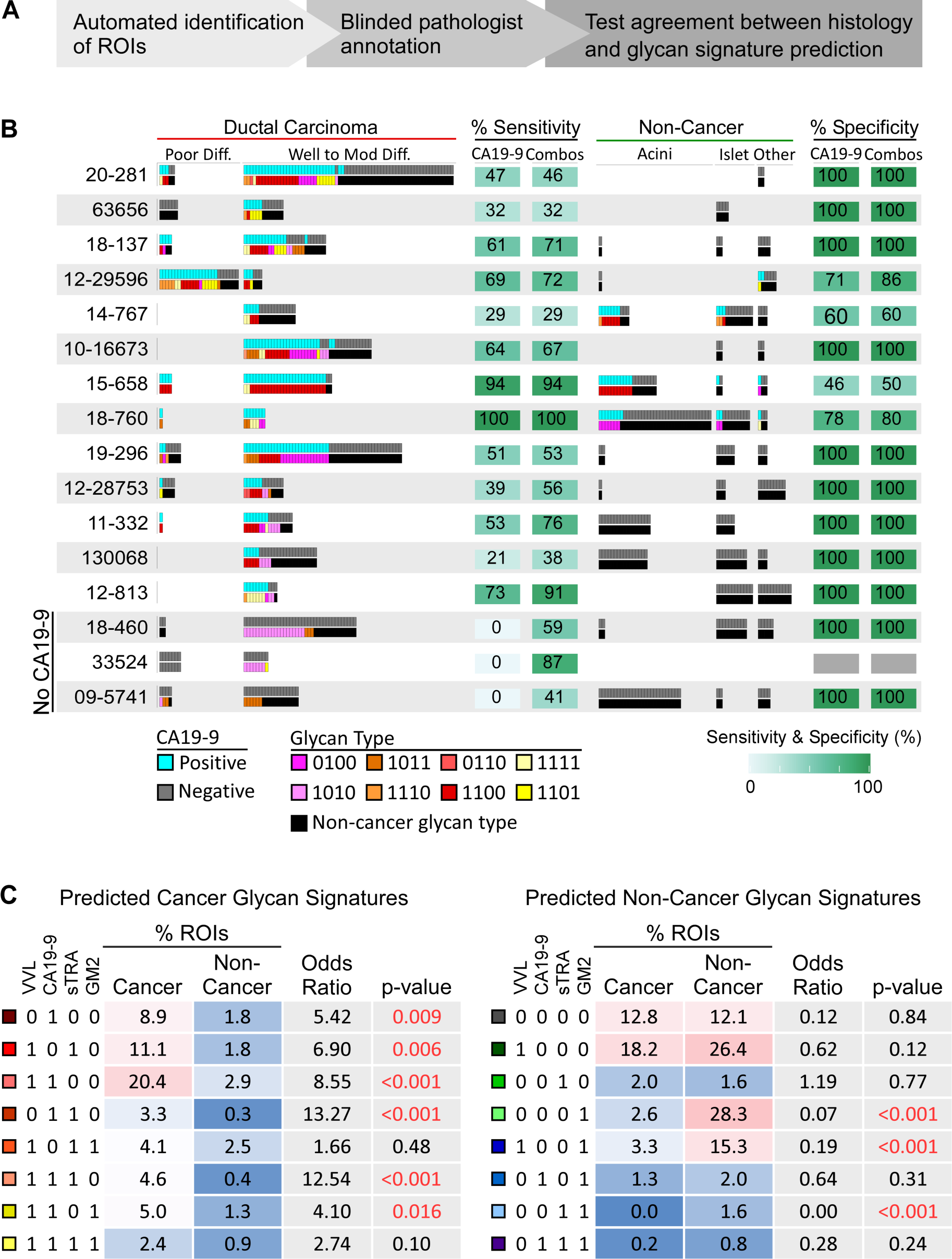
Test set validation of PDAC-associated glycan signatures. A) Test set design. B) Patterns of positivity of the putative cancer-associated glycan signatures compared to CA19-9 in individual ROIs in each specimen. C) Prevalence and association with cancer of the predicted cancer-associated (left) and non-cancer-associated glycan signatures.

Of the 8 putative cancer-associated glycan signatures determined from the training set, the test set demonstrated that 6 were significantly associated with PDAC with odds ratios (ORs) from 4.10 to 13.27 (p = 0.016 and p < 0.0001, respectively, inversion of bootstrap CI of OR (33)) (**Fig.3C**). Of the 8 predicted non-cancer glycan signatures, 3 were significantly associated with non-cancer with ORs of 0.19 or less (p < 0.0001, inversion of bootstrap CI of OR) (**Fig. 3C and Supplementary Table 6**). The ORs of the 2 non-significant predicted cancer signatures (1.66 and 2.74) were higher than all ORs of the predicted non-cancer signatures (**Fig. 3C**), showing that the direction and magnitude of all 8 of the predicted cancer-associated glycan signatures were aligned with the cancer ROIs.

We therefore investigated whether the use of the glycan signatures could improve the identification of cancer cells over the use of CA19-9 alone. We determined the number of ROIs that were positive for any of the 8 predicted cancer glycan signatures or for any of the 8 predicted non-cancer glycan signatures and compared these numbers to the pathologist-annotated histology of the ROIs. We made the same comparison using the numbers of ROIs positive or negative for CA19-9 alone. A significantly higher percentage of cancer ROIs were positive for the predicted cancer glycan signatures (59.7%, CI 51.7-69.3) than for CA19-9 alone (46%, CI 32.2-52, p < 0.0001, inversion of bootstrap CI for difference in sensitivity (33)), and an equivalent percentage of non-cancer ROIs were positive for the non-cancer glycan signatures (89.9%, CI 81.2 - 96.6) as were negative for CA19-9 alone (92.6%, CI 83.4 - 99.4, p = 0.24, inversion of bootstrap CI for difference in specificity) (**Supplementary Table 7**). We made similar observations in comparisons with sTRA and the other individual glycans (**Supplementary Table 7**). Thus, these analyses validated the associations with cancer of the 8 glycan signatures identified in the training set and supported the use of combinations of assays or hybrid assays to definitively identify these glycan signatures in tumor tissue.

### Segregated Distribution Among Spatial Clusters and Tumors of the Glycan-Defined Subpopulations

Previous research showed that PDAC subpopulations tend to segregate into spatially defined clusters within tumors, and that PDAC tumors variously contain either a single dominant subpopulation of PDAC cells or a mixture of subpopulations (**Fig. 4A**) (7, 8). Such spatial segregation is indicative of clonal outgrowths of cancer cells with diverging trajectories of DNA mutations (34, 35). The images of selected cancer ROIs suggested clonal outgrowth patterns with a general preponderance of one or two glycan-defined subpopulations, rather than a random mix (**Fig. 4B**). We therefore asked whether the spatial clusters of cancer cells within tumors were relatively homogeneously predominated by one or two cell subpopulations, or whether the cancer cell clusters contained random mixtures of cell subpopulations.

**Figure 4.**
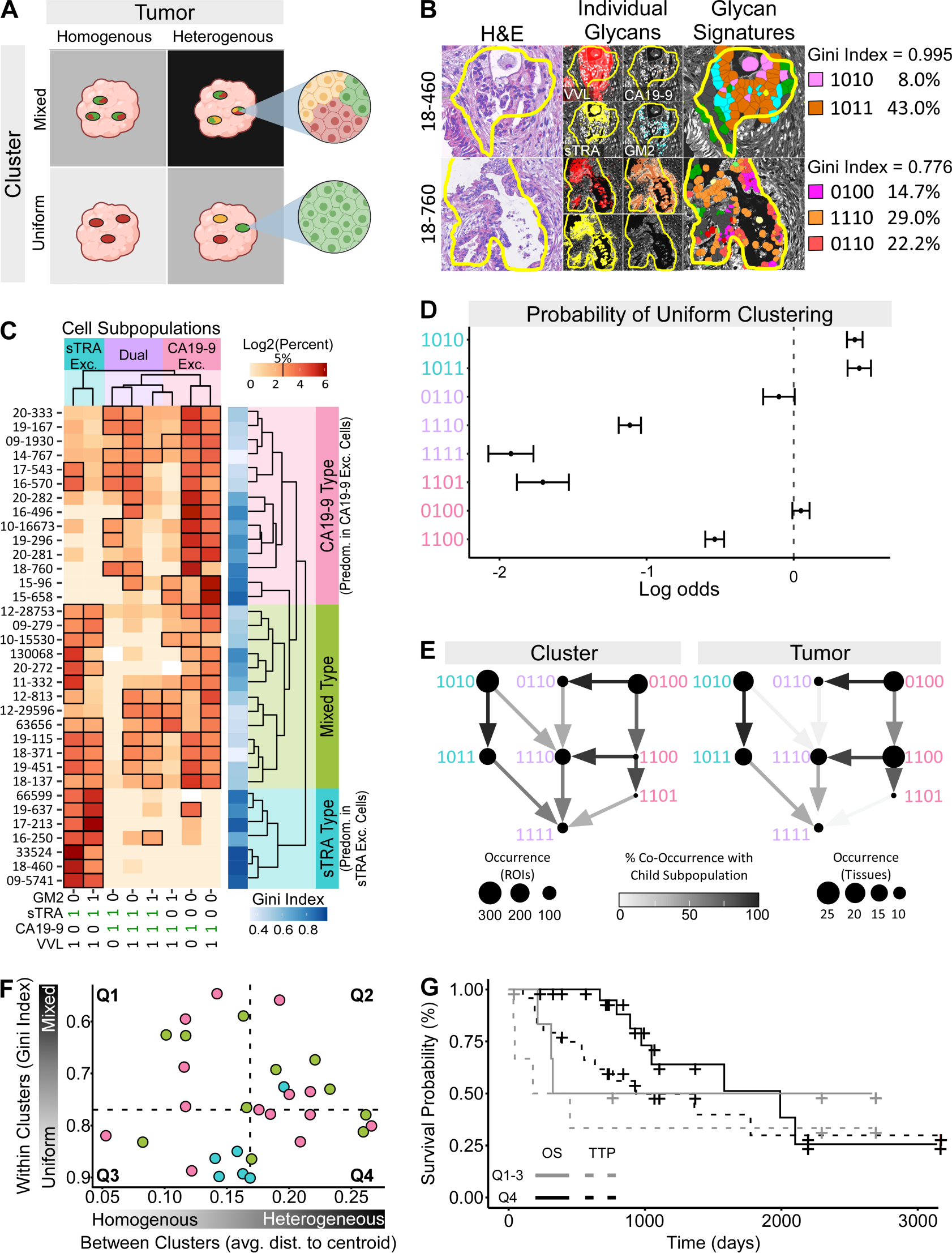
Unequal distribution of glycan-defined subpopulations among spatial clusters and tumors. A) Cluster and tumor heterogeneity. Clusters may be classified as uniform or mixed, and tumors may be classified as heterogeneous or heterogeneous. B) Two examples of within-cluster subpopulation purity, or uniformity, in cancer-cell clusters. C) Tumor type classifications based on glycan-signature type—CA19-9 exclusive, sTRA exclusive, or dual expressing— present in the tumor. D) Probability of association with pure, or uniform, clusters. E) Apparent lineages of glycan-defined PDAC cell subpopulations in the cell clusters and tumors. F) Relationship between average subpopulations purity of the clusters and the average centroid differences between clusters within each tumor. Each data point is a tumor, color coded by tumor type from panel C. The dashed lines are the median values in each axis. G) Kaplan-Meijer plot of PDAC patients in the different groups defined in panel F.

Using the whole set of 34 PDAC tumor samples, we identified regions within each tumor with the spatially clustered cancer-associated glycan signatures using the method described earlier (21) and determined the glycan signatures of all cells within the clusters (average 30,740 cells per tumor, total 1,045,142 cells). When we examined the spread of the percentages of each glycan-defined subpopulation across the tumors, we observed that the tumors clustered into three distinct groups of intratumoral distributions of glycan-defined subpopulations (**Fig. 4C**). One group of tumors contained mainly the PDAC cell subpopulations displaying sTRA in the absence of CA19-9 (codes 1010 and 1011, sTRA exclusive cells), and another group of tumors predominantly contained the subpopulations displaying CA19-9 in the absence of sTRA (codes 1100, 1101, and 0100, CA19-9 exclusive cells). A third group of tumors featured a heterogenous mix of subpopulations, including both the sTRA exclusive and the CA19-9 exclusive cells, as well as the dual expressing cells that display both CA19-9 and sTRA (codes 0110, 1110, and 1111). The Gini index of distribution inequality was divergent between the tumor groups, ranging from average 0.86 in the sTRA type tumors, to 0.63 in the CA19-9 type tumors, and to 0.54 in the mixed type tumors (**Fig. 4C**) (where 1 is completely homogeneous). Such a range between homogeneous and heterogeneous intratumoral distributions in cancer cell subpopulations mirrors the range observed with expression-defined subpopulations (7).

We then asked whether the glycan-defined PDAC cell subpopulations were equally or unequally distributed within the spatial clusters of cancer cells. Using the pathologist-annotated ROIs with PDAC histology (total 2,135 ROIs, average 122 per tumor), we found that the clusters, like the tumors, have distributions that range from relatively uniform (containing a single, predominant subpopulation) to mixed (**Supplementary Figure 5**). We then asked whether only certain glycan-defined cell subpopulations were found in the uniform clusters. We used a mixed effects model to determine the probability of any glycan-defined cell subpopulation appearing within a relatively uniform spatial cluster (Gini index ≥ 0.8) versus within a mixed spatial cluster (Gini index < 0.08) (**Fig. 4D**). We found that the sTRA exclusive (codes 1010 and 1011) and one of the CA19-9 exclusive (code 0100) subpopulations were significantly associated with uniform clusters, but that none of the dual expressing cell subpopulations were associated with uniform clusters (**Fig. 4D**).

Classical and basal-like PDAC subpopulations as defined by gene expression have patterns of homogeneity and heterogeneity within spatial clusters and tumors (7) that are similar to those observed here. These patterns suggested a lineage model of cancer progression from the classical to the mixed classical/basal-like to the basal-like subpopulations. We therefore asked whether our data are consistent with a lineage model of progression among the glycan-defined PDAC cell subpopulations. We determined the apparent lineages of the glycan-defined PDAC subpopulations using a network graph of their co-occurrences within spatial clusters (**Fig. 4E**). The relationships indicated that the sTRA exclusive and CA19-9 exclusive glycan signatures (codes 0100 and 1010, respectively) are independent of each other and rarely intermixed within clusters. The signatures that are positive for additional glycans, VVL and GM2, were subsets of the CA19-9 exclusive and sTRA exclusive cell subpopulations and also rarely intermixed in the same clusters (for example, code 1101 with code 1011). In addition, when we determined the apparent lineages at the tumor level using the percent of cells displaying each glycan signature within each tumor, we observed identical relationships (**Fig. 4E**). Thus, within this sample set, PDAC tumors occur in defined types that are either predominant in a single glycan-defined cell subpopulation or that have several subpopulations evenly intermixed among them. The sTRA exclusive and CA19-9 exclusive cells segregated into separate spatial clusters and tumors, consistent with a lineage model of progression from the sTRA exclusive and CA19-9 exclusive subpopulations to the dual expressing subpopulations.

If the glycan signatures identify subpopulations that align with clonal outgrowths of PDAC cells, one also would expect consistency with previous analyses suggesting that heterogeneity in clonal outgrowths of cancer cells is associated with poor outcome (36). We accordingly asked whether any tumors contain different kinds of pure PDAC cell clusters—postulated to be clonal outgrowths of different PDAC subpopulations—together in the same tumor, as modeled in **Figure 4A**. With this model, there are four classes of tumors: homogenous-uniform, homogenous-mixed, heterogeneous-uniform, and heterogeneous-mixed. For each tumor, we calculated both the average within-cluster uniformity (Gini index) and the between-cluster variance (average distance to centroid, **Supplementary Figure 5**). We found that a group of 8 tumors had both higher than median differences between the PDAC cell clusters and higher than median average purity within the clusters (**Fig. 4F**). We saw no general relationship between within-cluster homogeneity and between-cluster heterogeneity or with the tumor types defined in Figure 4C, indicating a broad range of characteristics among PDAC tumors. Based on the prediction that outgrowths of different clonal trajectories are predicted to be associated with poor outcomes (37), we compared survival probabilities between the patients with heterogenous-uniform tumors and the tumors in the other three groups (**Fig. 4G**). The time to 50% survival is shorter in the heterogeneous-uniform group (324 days vs 1991 days, respectively), and the 2-year survival rate is lower (50% vs 95%, respectively). In addition, survival after progression was significantly shorter in the heterogeneous-uniform group relative to the other three groups (average 197 vs. 470 days, respectively, p=0.048, Wilcoxon rank sum test). Overall survival was not significantly different between the groups (log-rank test), potentially owing to a small sample size, but the substantial difference in 50% and 2-year survival uniquely in the heterogeneous-uniform group suggests a link between clonal outgrowths of different PDAC subpopulations and poor survival.

### In-Vitro Assays for Cancer-Associated Glycan Signatures Enable Plasma-Based Identification of Different PDAC Tumor Types

We next asked whether the aberrant glycan signatures used in tumor tissue to distinguish between cancer cell subpopulations and tumor types also could be used in plasma samples in an analogous way. Different glycans displayed on the same cells can be released together on proteins or cell fragments such as extracellular vesicles (38) (**Fig. 5A**). A standard sandwich immunoassay captures and detects the same antigen, as in the CA19-9 assay. In contrast, a hybrid sandwich immunoassay uses different lectins or antibodies for capture and detection to bind and quantify the glycans present in a sample (**Fig. 5B**) (22, 39, 40). Such assays could provide improved cancer specificity by detecting glycans that typically are not co-produced by non-cancer cells, such as by capturing the CA19-9 antigen and detecting the sTRA antigen (referred to as the CA19-9.sTRA assay) (**Fig. 5B**).

**Figure 5.**
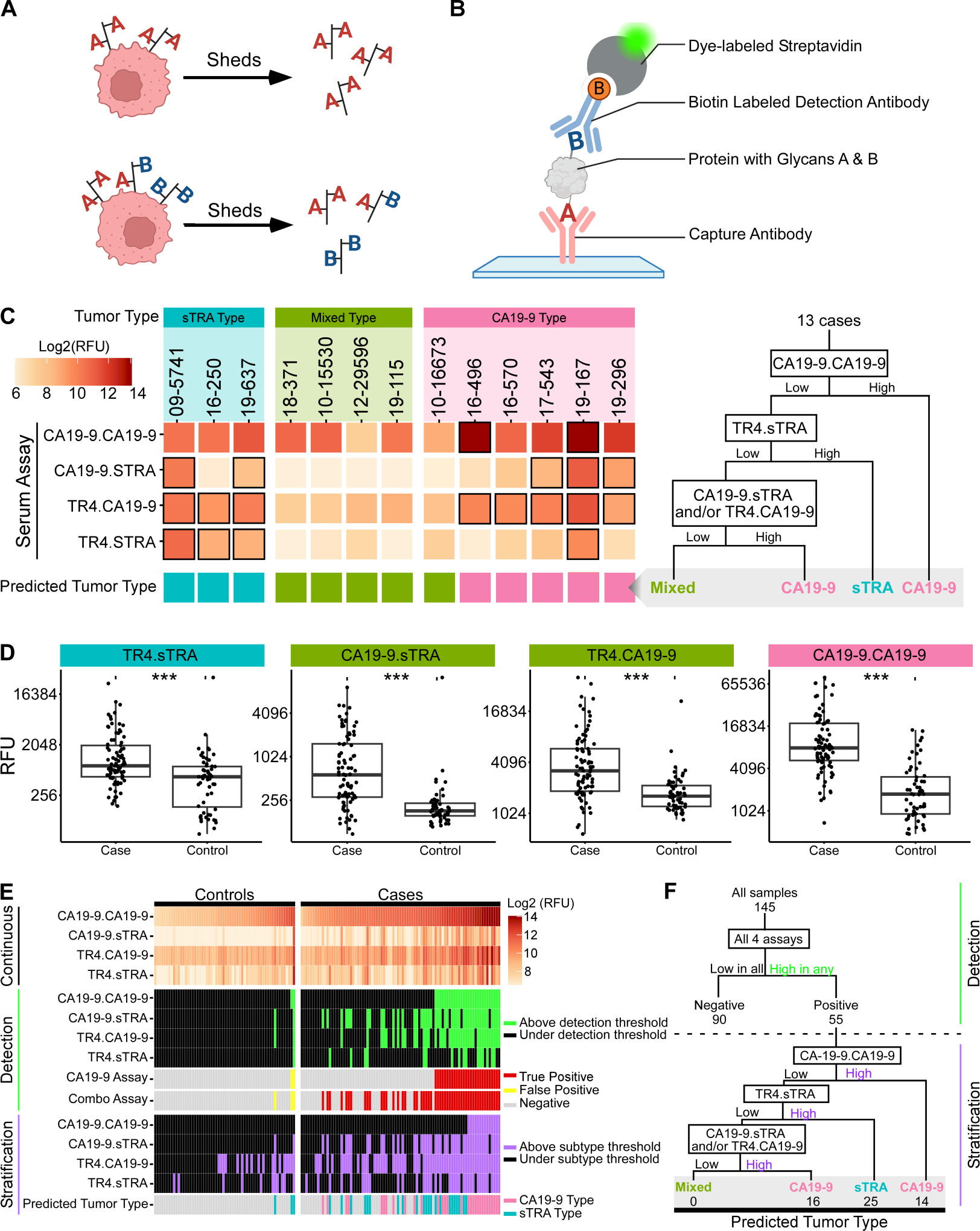
Complementary detection of tumors with divergent PDAC subpopulations using plasma assays. A) Model of co-secretion of glycans. B) Hybrid glycan sandwich assays. C) Correspondence of combined plasma assays to tumor types. D) Cancer-associated relative fluorescent unit (RFU) elevations in individual assays. E) Complementary detection of PDACs. F) Classification and stratification decision tree.

We first tested whether PDAC cancer cell lines release glycan-containing components that are detectable by *in-vitro* assays using conditioned media (**Supplementary Figure 6**). We used 24 different cell lines with a variety of classical or basal-like gene expression traits and a variety of glycan production traits (20). We cultured the cell lines on glass microscope slides divided into multiple wells per slide and processed these slides through our multiplexed glycan immunofluorescence and analysis pipeline. We confirmed the accuracy of the immunofluorescent glycan detection by comparisons with Western blots for the glycans (**Supplementary Figure 6**) and with chemiluminescence detection of the glycans on cells grown in plastic microtiter plates (not shown). At the cellular level, some cell lines were predominant in cells displaying the sTRA exclusive glycan signatures, others were predominant in cell displaying the CA19-9 exclusive glycan signatures, and others included cells displaying the dual expressing glycan signatures (**Supplementary Figure 6**). These distinct groups mirror those in the primary specimens but with lower heterogeneity, as expected in cell lines that potentially arise from a single cancer clone. We tested hybrid glycan assays using all combinations of capture and detection antibodies and lectins (16 assays total, substituting anti-Tn antibody for VVL as capture antibody). The conditioned media levels of each glycan significantly correlated across the cell lines with the corresponding percentages of cells with each glycan, and the relative levels between the hybrid glycan assays generally reflected the relative levels between the cellular glycan-defined subpopulations (**Supplementary Figure 6**), indicating that that media concentrations reflected the area covered by glycan-defined subpopulations on the slides. Thus, these data confirmed that the cell lines release these glycans into the media and that the *in-vitro* assays accurately capture and detect the glycans.

We then asked whether these hybrid glycan sandwich assays could detect glycans in PDAC patient derived blood samples that would correctly identify glycan-defined PDAC cell subpopulations within their matched tumor samples. We analyzed 13 blood samples (plasma and serum) that had a patient-matched tumor specimen in the above analysis. The patients with CA19-9 type tumors had the highest RFU values of secreted CA19-9 (determined using the CA19-9.CA19-9 assay), the patients with sTRA type tumors had the highest RFU values of secreted sTRA (determined using the TR4.sTRA assay), and the patients with mixed type tumors had low RFU values levels across all 4 assays (including CA19-9.sTRA and TR4.sTRA) (**Fig. 5C**). Using the measurements from all 4 assays, a decision tree with subtyping thresholds for each assay (**Supplementary Table 8**) correctly classified each blood sample by the type of the matched tumor in all but one case (**Fig. 5C**). Thus, using the combined measurements from the 4 assays, the blood sample measurements provided an indirect identification of the different types of PDAC tumors.

To determine whether our hybrid glycan sandwich assays could distinguish between plasma from PDAC patients and plasma from control subjects, we analyzed plasma samples from 85 PDAC patients and 60 control subjects (**Supplementary Table 9**). The PDAC subjects had resectable cancer (stage I/II) in 59/85 (69%) cases and stage III/IV cancer in 18/85 (21%) cases (8/85 (9.4%) not available), and the control subjects included healthy individuals as well as patients with benign conditions requiring differential diagnosis, such as chronic pancreatitis and benign biliary obstruction. We observed that the measurements from each of the 4 assays was significantly higher in the cancer cases than in the controls (p < 0.05, Wilcoxon rank sum test) (**Fig. 5D**). No significant associations were detected between any of the assays and sex, age, or type of sample (serum vs plasma) (Fisher Exact test).

Therefore, for detection, we combined the assays to produce a single test, in which a sample was classified as a case if above the high-specificity cutoff in any assay (**Fig. 5E**), as demonstrated previously (17, 41). To estimate the sensitivity (true positive detection) of the combined assays and CA19-9 by itself, we performed 1000-fold resampling of the data with threshold selection in each sample based on achieving 95% specificity (true negative detection) in the controls. The estimated sensitivity was significantly greater for the combined assays (62%, 53 of 85 PDACs) relative to CA19-9 by itself (29%, 25 of 85 PDACs) (p = 0.042, inversion of bootstrap CI of the difference), with equivalent specificity for the combined assays (95%, 57 of 60 controls) and CA19-9 by itself (100%, 60 of 60 controls) (p = 0.380, inversion of bootstrap CI of the difference). Each of the additional assays besides CA19-9 showed unique elevations among the PDAC patients, indicating complementary contributions to the improved detection performance (**Fig. 5E**). This significant improvement in sensitivity over CA19-9 is consistent with the ability to detect both the tumors with predominantly CA19-9 exclusive subpopulations and the tumors with predominantly sTRA exclusive subpopulations and signifies future enhanced value for PDAC detection.

The ability to detect PDACs with different predominant subpopulations not only has value for earlier detection but also for the stratification of patients based on their tumor characteristics. We therefore simulated a two-stage process of detection followed by stratification, in which we used the blood measurements first to detect PDAC and then to predict the type of PDAC tumor (**Fig. 5E**). Among the patients that were classified as PDAC by the combined blood assays, we predicted the PDAC tumor type using the cutoffs and decision tree developed above (**Fig. 5E**). Of the 55 subjects classified as PDAC, 30 were predicted to have the CA19-9 type tumor and 25 were predicted to have the sTRA type tumor. We were not able to detect or predict tumors of the mixed type due to the lack of high RFU values of the individual blood sample assays for that tumor type. Thus, these results indicated that the combined hybrid-glycan assays provide a means to improve PDAC detection and stratification using a blood test to identify PDACs across the spectrum of tumor types.

## Discussion

In the present report, we have made three observations. First, PDAC cancer cell subpopulations display 2-4 glycan signatures of cancer that are rarely found on non-cancer cells and that may be detected using our novel multiplexed glycan immunofluorescence pipeline. Second, using our novel pipeline, we have identified three broad PDAC cancer cell subpopulations: those defined by CA19-9 alone in the absence of sTRA (CA19-9 exclusive), those defined by sTRA alone in the absence of CA19-9 (sTRA exclusive), or those defined by both CA19-9 and sTRA together (dual expressing). Similarly, at the tumor level, we observed three tumor types that are defined broadly by the presence of predominantly CA19-9 exclusive cells (CA19-9 type), the presence of predominantly sTRA exclusive cells (sTRA type), or the presence of both subpopulations (mixed type). Third, using our novel hybrid glycan immunoassays to detect pairs of membrane-associated glycans, we found in patient matched tumor tissue and blood samples that the relative abundances of the detected serum glycans were consistent with the same glycans at the tumor tissue levels. Therefore, our novel hybrid glycan immunoassays identified both the CA19-9 type and the sTRA type tumors and showed improved sensitivity and specificity when compared with CA19-9 alone. Because of these technical and conceptual innovations, this methodology could enable the clinical development of a simple blood test that would improve early detection of patients with early PDAC.

Previous approaches to differentiating between PDAC cancer cell subpopulations have been based on multi-gene and multi-protein expression profiles in tumor tissue (2–4, 7). This approach has been foundational for defining the classical, basal-like, and additional PDAC subpopulations, but it also is clinically limited because previously investigated RNA and protein biomarkers, such as GATA6 to identify classical PDAC or KRT17 to identify basal-like PDAC, are not detectable in the peripheral blood. Glycans have the advantage of being detectable, in many cases, on both the cell surfaces and in the blood circulation. Previous studies of individual glycans, such as sialic acids (42), O-linked glycans (43, 44), N-linked glycans examined by MALDI imaging (45), sialyl Lewis X (39, 46), and others (47), supported the hypothesis that PDAC cancer cells display altered glycan signatures when compared with relevant non-cancer controls. These earlier studies did not employ multiplexing methods or the new image analysis approaches that allow for broad, statistical analysis across multiple cell types and tumors. Our approach, however, allowed for robust, multiplexed statistical analysis due to the automated signal thresholding of SignalFinder. Binary coding then enabled the tracking and analysis of the glycan signatures observed in specific cell types. The results show proof of concept that glycan signatures may be used as biomarkers for PDAC subpopulations and, in turn, early PDAC detection.

The glycan-defined PDAC subpopulations that we identified segregate in a similar way to the canonical, expression-based classical and basal-like PDAC subpopulations. For example, the cells with CA19-9 exclusive signatures aligned with well differentiated PDAC (**Supplementary Figure 3**) and, in a previous study of cell lines (20), with epithelial morphology and the classical gene-expression signature, which is consistent with the canonical classical subpopulation. Similarly, the cells with sTRA exclusive signatures aligned with poorly differentiated PDAC (**Supplementary Figure 3**) and previously with mixed epithelial/mesenchymal morphology and the basal-like gene expression signature (20), which is consistent with the canonical basal-like subpopulation. Further, the dual expressing cells which express both CA19-9 and sTRA may correspond to the dual classical-basal cell subpopulation as they both theoretically derive from the same canonical subpopulations. An association between cell subpopulations and distinct glycan signatures concords with the regulation of glycan production, which involves the expression of glycosyltransferases, the metabolism of monosaccharides, and the routing of substrates (13–15). Distinct PDAC cancer cell subpopulations develop through outgrowths from progenitor populations (40), changes in the microenvironment (41), or alterations in epigenetic drivers (42), potentially bringing substantial changes to each of these processes and, as a result, remodeling of the cell-surface and secreted glycans. Thus, the glycan signatures of the distinct PDAC subpopulations may be unique to the underlying biological differences between the subpopulations.

Our glycan binary codes determined from tumors were translatable to a blood test using a set of novel hybrid glycan assays. We indirectly identified from the blood assays the type of PDAC subpopulations that were predominant in the corresponding tumors. This was done by a straightforward assessment of positivity or negativity in 4 hybrid-glycan assays—one detecting only CA19-9, one detecting only sTRA, and two detecting linked CA19-9 and sTRA. This test has the advantages of requiring only 4 standard immunoassays, minimal sample volume, and a straightforward decision algorithm. The ability to detect the different PDAC tumor types led to the detection of 62% of the PDACs, relative to 33% by CA19-9 alone using cutoffs that gave low false-positive detection (5% and 3%, respectively). Such substantial improvements upon CA19-9 using a practical assay could be immediately valuable, given that CA19-9 already is being evaluated as part of algorithms to enhance PDAC surveillance among elevated-prevalence populations (48). Thus, with minor modifications, this methodology could be an efficient and economical approach to improving early PDAC detection.

Blood assays for PDAC subpopulation-associated glycan signatures also could have value in clinical applications beyond early PDAC detection. In cell-culture (2), organoid (49), and retrospective patient studies (2–4), the tumors that are predominant in the classical and basal-like subpopulations show differential responses to treatments, but, owing to the lack of blood biomarkers, these findings are not tested in human studies to track responses in drug trials. Glycan signatures that correspond to the gene-expression defined subpopulations would provide a means to translate the cell line and organoid findings to human studies. Glycan signatures could also provide additional information based on their potential functional roles in cancer progression. The cell-surface glycome maintains the integrity of multicellular organization, potentially enables migration and extravasation, mediates metastatic colonization, modulates immune recognition, and regulates the promotion and suppression of inflammation (50–53). CA19-9 and related Lewis-family glycans could promote metastatic cell seeding through interactions with selectin-family receptors (54), and sialylated glycans like sTRA interact with siglec receptors that lead to immunosuppression (42). The distinct glycan-defined PDAC subpopulations could thus have differing propensities to metastasize or have differing responses to immunotherapies and other treatments. Consequently, the ability to detect and distinguish the PDAC subpopulations by glycan signatures may enable improved patient stratification and targeted therapy.

A limitation of the current study is that the matched set of 13 tumor and 13 blood samples may not be fully representative of the broader PDAC population. The study also used 1 tumor block per case, which in some samples represent a minority fraction of the tumor and therefore may not provide an accurate representation of the whole tumor. Larger patient-matched sets will be required to establish the accuracy of the blood assays for tumor type prediction. Larger cohorts also will be needed to determine the associations between intra-tumoral heterogeneity in PDAC subpopulations and outcomes. The current study also revealed that the current set of 4 hybrid glycan assays detect mainly the CA19-9-type and sTRA-type tumors but not the heterogeneous tumors. As a result, the current blood biomarkers identify only the tumors that are CA19-9 type or sTRA type. However, the present work shows proof of concept that we can develop hybrid-glycan biomarker assays for the mixed type tumors through additional glycans that potentially improve the detection of mixed type tumors. In addition, the performance of the combined assays for the discrimination of early PDAC from controls will need to be determined in larger case/control studies and eventually in prospective studies to fully evaluate applicability to PDAC surveillance. In the current study, the sensitivity and specificity of CA19-9 and sTRA used in combination were consistent with previous observations (17, 20, 23, 41), supporting the accuracy of the findings.

Here, for the first time we show that we can identify the presence of specific PDAC cancer cell subpopulations in tumors using a simple blood test and that this improves the overall identification of incipient PDAC cases from controls. We achieve this goal through the application of multiplexed glycan immunofluorescence to primary PDAC tumors and the detection of tumor-released pairs of glycans in the blood plama with a novel hybrid glycan sandwich assay. A blood test to detect tumors with divergent levels of distinct PDAC subpopulations could enhance PDAC identification in surveillance for PDAC. In addition, based on the ability to identify tumor PDAC subpopulations using blood samples, one can translate cell line, organoid, and model-system studies of metastatic potential and therapeutic or radiation susceptibility to human studies that track the time course of subpopulation-specific responses to treatment regimens. This in turn could inform how PDAC is identified, stratified, and treated in the clinic.

## Methods

### Human Specimens

The tissue samples were collected from surgical resections of PDAC patients at Duke University Medical Center, University of Pittsburgh Medical Center, Trinity Health Grand Rapids, or the University Health Network, Toronto. All samples were formalin fixed and paraffin embedded. The tissue sections used for this study were from tissue blocks not needed for patient evaluation and patient outcomes were recorded for at least 3 years.

Human blood plasma or serum specimens were assembled through the clinical centers at Duke University Medical Center, University of Pittsburgh Medical Center, Trinity Health Grand Rapids, or MD Anderson Cancer Center. The samples were collected prior to any cancer treatment and were processed under a standard operating procedure approved by the Early Detection Research Network (55). All plasma samples used EDTA anti-coagulant and were frozen at -80° C within 3 hours of collection. The aliquots used in the study underwent no more than 3 freeze/thaw cycles prior to use.

### Sex As a Biological Variable

The study included statistically equivalent representation of male and female subjects. There were no significant differences between sex in the tumor fractions of any glycan signature in PDAC cells (Wilcoxon Rank Sum test). There was no significant difference in sensitivity or specificity between sex for the individual or combined blood assays (Fisher Exact test).

### Multiplexed Immunofluorescence

We performed immunofluorescence on 5 µm thick sections cut from formalin-fixed, paraffin-embedded (FFPE) blocks. Deparaffinization performed by CitriSolv Hybrid (Decon Labs, King of Prussia, PA) that contained d-limonene and isopropyl cumene and rehydrated through an ethanol gradient of 100%, 95%, and 70% followed by washing with PBS. Following rehydration, antigen retrieval was achieved through incubating slides in citrate buffer at 100 °C for 20 minutes. Slides were blocked with phosphate-buffered saline with 0.05% Tween-20 (PBST 0.05) and 3% bovine plasma albumin (BSA) for 1 hour at room temperature (RT). Tissue staining was manual and performed in multiple rounds. The primary antibodies and lectins (**Supplementary Table 1**) were labeled for immunofluorescence with sulfo-cyanine-5 (Cy5) NHS ester or sulfo-cyanine-3 (Cy3) NHS ester (Lumiprobe, MD), according to instructions from the provider. After dialysis to remove unreacted conjugate, a Cy5-labeled antibody or lectin and a Cy3-labeled antibody or lectin were mixed into the same solution of PBST 0.05 with 3% BSA to a final concentration of 10 µg/mL. Slides were incubated overnight with this solution at 4 °C in a humidified chamber.

The following day, the solutions were decanted, and the slides were washed twice in PBST 0.05 and once in 1X PBS, each time for 3 minutes. The slides were dried via blotting and incubated with DAPI (AnaSpec, Fremont, CA) at 10 µg/mL in 1X PBS for 15 minutes at RT. Two five-minute washes were performed in 1X PBS, and then slides were cover-slipped and scanned using a fluorescent microscope (AxioScan.Z1, Zeiss, Oberkochen, Germany). We next quenched the fluorescence using 6% H2O2 in 250 mM sodium bicarbonate (pH 9.5-10) and performed another round of immunofluorescence using two different antibodies or lectins, using the order of VVL and GM2 in round 1, CA19-9 and TRA-1-60 in round 2, and TRA-1-60 and GSL-II in round 3. Prior to the second round of detection with the TRA-1-60 antibody, we treated the slides with sialidase to remove terminal sialic acids. For this step, the slides were incubated with a 1:200 dilution (from a 50,000 U/mL stock) of α2-3,6,8 neuraminidase in 5 mM CaCl2, 50mM pH 5.5 sodium acetate overnight at 37 °C.

In each round of fluorescence imaging, the microscope collected three images at each field-of-view, each image corresponding to the emission maxima of DAPI, Cy3, and Cy5. The three images were saved as independent, stacked layers in a single file. Following all fluorescence imaging, the slides were stained with hematoxylin and eosin (H&E) using a standard protocol and digitally imaged using the Aperio ScanScope (Leica).

### Image Analysis

All fluorescence images were initially processed using SignalFinder. SignalFinder automatically creates a map of the locations of pixels containing signal in each layer of the multiplexed immunofluorescence as an ome.tiff. This map was aligned to the H&E image based on the signals from cell nuclei using the Warpy extension in ImageJ (31, 32). We used automated cell nuclei detection followed by a 10 µm expansion to estimate cell borders and then quantified the fractional area of each cell that was positive for each image layer. Each image layer corresponded to a single glycan, yielding a set of 6 fractional-positive values for each cell.

### Bioinformatics Methods

We trained a classification tree to distinguish the cancer from the non-cancer cells using the 6 fractional-positive values (CA19-9, TRA, GM2, VVL, sTRA, GSL-II) for each cell. The values from TRA and GSL-II had the lowest importance scores and did not affect classification accuracy when removed and were therefore not used in subsequent analyses (**Supplementary Figure 2**). A classification tree created from the remaining 4 glycans was used to select thresholds in each fractional-positive value to dichotomize each cell as positive or negative for each glycan. The thresholds were selected from the first split in the tree for each glycan, using trees from multiple initial selections of the training and validation sets in cross validation. We then used the dichotomized glycan data in a classification tree to identify signatures of glycans associated with cancer or non-cancer. Each terminal branch of the resulting tree was assigned a 4-digit binary code based on the sequence of positive or negative splits for each glycan and assigned a cancer or non-cancer classification based on the majority of cells in the branch.

The analysis of apparent tumor lineage was done through network graphs of glycan signature occurrence and co-occurrence, where occurrence was defined as a minimum of 5% of cells positive for the glycan signature within the cluster or tumor. Percent co-occurrence is the fraction of observations in the child node which also occur in the parent node. The network graph only considers stepwise accumulation of glycans.

### Immunoassays

The immunoassays were based on the method presented earlier (22, 39). Capture antibodies were deposited in arrays on microscope slides that were functionalized to enable covalent attachment (Z Biotech, 10401-2) using a microarray printer (Dispendix I.DOT-One). The capture antibodies anti-CA19-9, anti-TR4, anti-GM2, and anti-Tn (Supplementary Table 1) were printed in sodium borate printing buffer (pH 8.5) with 7.5% glycerol at 50 µg/mL. Individual arrays were separated with a 64-well gasket system (248865, Grace Bio-Labs, Bend, OR). We diluted the samples of human plasma or serum (8-fold) and cell line conditioned media (2-fold) into a buffer (1X PBS with 0.1% Tween-20, 0.1% Brij-35, species-specific blocking antibodies, and protease inhibitor) and incubated each sample on an antibody array (pre-blocked with 1% BSA) overnight. For sTRA detection arrays, incubation with the unlabeled TRA-1-60 antibody (to block native epitopes) was performed before sialidase treatment to expose the sialylated epitopes (22). We prepared α2-3 neuraminidase (P0728L, New England Biolabs, Ipswich, MA) at 250 U/mL in GlycoBuffer1 (50 mM sodium acetate, 5 mM CaCl_2_) and incubated the solution on the arrays for one hour or overnight at 37 °C. Next, we incubated all arrays with a biotinylated detection antibody or lectin (3 μg/mL in 1X PBS with 0.1% Tween-20 and 0.1% BSA) and subsequently with Cy5-conjugated streptavidin (43-4316, Invitrogen, Carlsbad, CA) (2 μg/mL in the same buffer as the primary antibody). The slides were scanned for fluorescence at 635 nm using a microarray scanner (Innopsys InnoScan 1100 AL). The quantification of fluorescence was performed using SignalFinder-Microarray (56).

### Statistical Analysis

Analysis of glycan signature differences between cell types (Figure 2D and Supplemental Figure 2B) was performed using the Wilcoxon Rank Sum test with stratification of ROIs by specimen identifier to correct for repeated measures. This was done using the “clusRank” R package (57). The p-values for this test were corrected using the false discovery rate (FDR) correction procedure as implemented in the “p.adjust” R function.

For estimation of biomarker performance, such as sensitivity, specificity, difference in sensitivity or specificity (Figure 5E), and odds ratio with cancer/non-cancer, using ROIs from PDAC specimens (Figure 3C), the 95% percentile bootstrap confidence intervals the estimator were constructed by 1000 bootstrap sampling of the PDAC specimens with replacement, and the p-values were calculated by inversion of the percentile bootstrap CI using the “boot.pval” R package.

Analysis of cell odds of belonging to different cell types (Supplemental Figure 2A) or clusters of different uniformity (Figure 4D) was calculated using a binomial mixed effects regression model with the logit (log-odds) link function as defined in the “glmer” function of the “lme4” package (58); we used the “0000” glycan signature as the reference group. Variance between clusters within a tumor was calculated as the average distance of cancer associated glycan signatures distributions of each cluster for a given tumor to the centroid (multivariate median). This was calculated using the “betadisper” function of the “vegan” R package.

### Software

We developed the SignalFinder software using MATLAB and C++. Cellular and spatial signal analysis was done in MATLAB and QuPath. We used Microsoft Excel and R for analyzing numerical output and the preparation of graphs, BioRender for the preparation of graphical cartoons, and Canvas XIV for the preparation of the composite figures.

### Study Approval

The tissue samples were collected under protocols approved by the Institutional Review Boards at Duke University Medical Center, University of Pittsburgh Medical Center, Trinity Health Grand Rapids, or the University Health Network, Toronto. The human blood plasma or serum specimens were collected by each site under the approval of the institutional review boards at Duke University Medical Center, University of Pittsburgh Medical Center, Trinity Health Grand Rapids, or MD Anderson Cancer Center. All subjects provided written, informed consent, and in accordance with an assurance filed with and approved by the U.S. Department of Health and Human Services.

### Data and Software Availability

The data derived from this study and the SignalFinder software are available upon request.

## Additional Information

## Author Contributions

Experimental design was performed by CG, BS, BB, ZK, and BH. Experiments and data collection were performed by CG, BS, and BB. Pathology review was performed by GH and CS. Data analysis and modeling AR, BB, ZK, and YH. Software development was performed by HLT and ZK. Specimens were provided by JM, SG, PA, AS, and RB, and reagents were provided by MB. Statistical analysis was performed by YH, AR, and ZK. Manuscript was written by BH, DB, BB, and ZK. BB, ZK, BH, and DB provided significant intellectual contributions. BB and ZK are co-first authors for their cumulative contributions throughout the project, with BB providing major intellectual contributions and ZK providing major analytical contributions. All authors read and approved the manuscript

## Acknowledgements

We thank the members of the Biorepository and Pathology Core (Van Andel Institute) for assistance with preparation and processing of the tumor and plasma/serum specimens and Kristin Gallik, PhD, of the Optical Imaging Core (Van Andel Institute) for assistance in acquisition of the multiplexed immunofluorescence images and for QuPath training and support. We thank Anirban Maitra, PhD, (MD Anderson Cancer Center) for providing plasma specimens.

## References

1. Siegel RL, et al. Cancer statistics, 2023. CA: A Cancer J Clin. 2023;73(1):17–48.

2. Collisson EA, et al. Subtypes of Pancreatic Ductal Adenocarcinoma and Their Differing Responses to Therapy. Nat Med. 2011;17(4):500–503.

3. Moffitt RA, et al. Virtual microdissection identifies distinct tumor- and stroma-specific subtypes of pancreatic ductal adenocarcinoma. Nat Genet. 2015;47(10):1168–1178.

4. Bailey P, et al. Genomic analyses identify molecular subtypes of pancreatic cancer. Nature. 2016;531(7592):47–52.

5. Chan-Seng-Yue M, et al. Transcription phenotypes of pancreatic cancer are driven by genomic events during tumor evolution. Nat Genet. 2020;52(2):231–240.

6. Hwang WL, et al. Single-nucleus and spatial transcriptome profiling of pancreatic cancer identifies multicellular dynamics associated with neoadjuvant treatment. Nat Genet. 2022;54(8):1178–1191.

7. Williams HL, et al. Spatially Resolved Single-Cell Assessment of Pancreatic Cancer Expression Subtypes Reveals Co-expressor Phenotypes and Extensive Intratumoral Heterogeneity. Cancer Res. 2022;83(3):441–455.

8. Juiz N, et al. Basal-like and classical cells coexist in pancreatic cancer revealed by single-cell analysis on biopsy-derived pancreatic cancer organoids from the classical subtype. Faseb J. 2020;34(9):12214–12228.

9. Carpenter ES, et al. KRT17High/CXCL8+ tumor cells display both classical and basal features and regulate myeloid infiltration in the pancreatic cancer microenvironment. Clin Cancer Res. 2023;OF1–OF17.

10. Topham JT, et al. Subtype-Discordant Pancreatic Ductal Adenocarcinoma Tumors Show Intermediate Clinical and Molecular Characteristics. Clin Cancer Res. 2021;27(1):150–157.

11. O’Kane GM, et al. GATA6 Expression Distinguishes Classical and Basal-like Subtypes in Advanced Pancreatic Cancer. Clin Cancer Res. 2020;26(18):4901–4910.

12. Aung KL, et al. Genomics-Driven Precision Medicine for Advanced Pancreatic Cancer - Early Results from the COMPASS Trial. Clin Cancer Res. 2017;24(6):1344–1354.

13. Araujo L, et al. Glycolysis and glutaminolysis cooperatively control T cell function by limiting metabolite supply to N-glycosylation. eLife. 2017;6:e21330.

14. Lucena MC, et al. Epithelial Mesenchymal Transition Induces Aberrant Glycosylation through Hexosamine Biosynthetic Pathway Activation. J Biol Chem. 2016;291(25):12917– 12929.

15. Rossi M, et al. PHGDH heterogeneity potentiates cancer cell dissemination and metastasis. Nature. 2022;605(7911):747–753.

16. McDowell CT, et al. Imaging Mass Spectrometry and Lectin Analysis of N-Linked Glycans in Carbohydrate Antigen–Defined Pancreatic Cancer Tissues. Mol Cell Proteomics. 2021;20:100012.

17. Staal B, et al. The sTRA Plasma Biomarker: Blinded Validation of Improved Accuracy over CA19-9 in Pancreatic Cancer Diagnosis. Clin Cancer Res. 2019;25(9):2745–2754.

18. Ballehaninna UK, Chamberlain RS. The clinical utility of serum CA 19-9 in the diagnosis, prognosis and management of pancreatic adenocarcinoma: An evidence based appraisal. J Gastrointest Oncol. 2011;3(2):105–19.

19. Barnett D, et al. The CA19-9 and Sialyl-TRA Antigens Define Separate Subpopulations of Pancreatic Cancer Cells. Sci Reports. 2017;7(1):4020.

20. Gao C, et al. Detection of Chemotherapy-resistant Pancreatic Cancer Using a Glycan Biomarker, sTRA. Clin Cancer Res. 2021;27(1):226–236.

21. Wisniewski L, et al. Heterogeneity of glycan biomarker clusters as an indicator of recurrence in pancreatic cancer. Frontiers Oncol. 2023;13:1135405.

22. Tang H, et al. Glycans Related to the CA19-9 Antigen Are Increased in Distinct Subsets of Pancreatic Cancers and Improve Diagnostic Accuracy Over CA19-9. Cell Mol Gastroenterology Hepatology. 2015;2(2):210–221.e15.

23. Barnett D, et al. The CA19-9 and Sialyl-TRA Antigens Define Separate Subpopulations of Pancreatic Cancer Cells. Sci Rep-uk. 2017;7(1):4020.

24. Klamer ZL, et al. CarboGrove: a resource of glycan-binding specificities through analyzed glycan-array datasets from all platforms. Glycobiology. 2022;32(8):679–690.

25. Springer GF. T and Tn, General Carcinoma Autoantigens. Science. 1984;224(4654):1198–91206.

26. Ju T, et al. Human Tumor Antigens Tn and Sialyl Tn Arise from Mutations in Cosmc. Cancer Res. 2008;68(6):1636–1646.

27. Sinha J, et al. A Gastric Glycoform of MUC5AC Is a Biomarker of Mucinous Cysts of the Pancreas. Plos One. 2016;11(12):e0167070.

28. Andrews PW, et al. Three Monoclonal Antibodies Defining Distinct Differentiation Antigens Associated with Different High Molecular Weight Polypeptides on the Surface of Human Embryonal Carcinoma Cells. Hybridoma. 1984;3(4):347–361.

29. Tan L, et al. A Targetable Pathway to Eliminate TRA-1-60+/TRA-1-81+ Chemoresistant Cancer Cells. J Mol Cell Biol. 2023;6(15):mjad039.

30. Sasaki N, et al. Ganglioside GM2, highly expressed in the MIA PaCa-2 pancreatic ductal adenocarcinoma cell line, is correlated with growth, invasion, and advanced stage. Sci Rep-uk. 2019;9(1):19369.

31. Schindelin J, et al. Fiji: an open-source platform for biological-image analysis. Nat Methods. 2012;9(7):676–682.

32. Chiaruttini N, et al. An Open-Source Whole Slide Image Registration Workflow at Cellular Precision Using Fiji, QuPath and Elastix. Front Comput Sci. 2022;3:780026.

33. Thulin M. Modern Statistics with R. EOS Chasma Press; 2021.

34. Malinova A, et al. Cell Lineage Infidelity in PDAC Progression and Therapy Resistance. Front Cell Dev Biol. 2021;9:795251.

35. Braxton AM, et al. 3D genomic mapping reveals multifocality of human pancreatic precancers. Nature. 2024;1–9.

36. Krieger TG, et al. Single-cell analysis of patient-derived PDAC organoids reveals cell state heterogeneity and a conserved developmental hierarchy. Nat Commun. 2021;12(1):5826.

37. Dagogo-Jack I, Shaw AT. Tumour heterogeneity and resistance to cancer therapies. Nat Rev Clin Oncol. 2018;15(2):81–94.

38. Wu L, Gao C. Comprehensive Overview the Role of Glycosylation of Extracellular Vesicles in Cancers. ACS Omega. 2023;8(50):47380–47392.

39. Tang H, et al. Glycan Motif Profiling Reveals Plasma Sialyl-Lewis X Elevations in Pancreatic Cancers That Are Negative for Sialyl-Lewis A. Mol Cell Proteomics. 2015;14(5):1323–1333.

40. Chen S, et al. Multiplexed analysis of glycan variation on native proteins captured by antibody microarrays. Nat Methods. 2007;4(5):437–444.

41. Haab B, et al. A Rigorous Multi-Laboratory Study of Known PDAC Biomarkers Identifies Increased Sensitivity and Specificity Over CA19-9 Alone. bioRxiv. 2024;2024.05.22.595399.

42. Rodriguez E, et al. Sialic acids in pancreatic cancer cells drive tumour-associated macrophage differentiation via the Siglec receptors Siglec-7 and Siglec-9. Nat Commun. 2021;12(1):1270.

43. Freitas D, et al. O-glycans truncation modulates gastric cancer cell signaling and transcription leading to a more aggressive phenotype. EBioMedicine. 2019;40:349–362.

44. Chugh S, et al. Disruption of C1galt1 Gene Promotes Development and Metastasis of Pancreatic Adenocarcinomas in Mice. Gastroenterology. 2018;155(5):1608–1624.

45. McDowell CT, et al. Applications and continued evolution of glycan imaging mass spectrometry. Mass Spectrom Rev. 2023;42(2):674–705.

46. Pour PM, et al. Expression of blood group-related antigens ABH, Lewis A, Lewis B, Lewis X, Lewis Y, and CA 19–9 in pancreatic cancer cells in comparison with the patient’s blood group type. Cancer Res. 1988;48(19):5422–6.

47. Munkley J. The glycosylation landscape of pancreatic cancer. Oncol Lett. 2019;17(3):2569– 2575.

48. Fahrmann JF, et al. Lead-Time Trajectory of CA19-9 as an Anchor Marker for Pancreatic Cancer Early Detection. Gastroenterology. 2021;160(4):1373–1383.e6.

49. Tiriac H, et al. Organoid profiling identifies common responders to chemotherapy in pancreatic cancer. Cancer Discov. 2018;8(9):1112–1129.

50. McKitrick TR, et al. The Crossroads of Glycoscience, Infection, and Immunology. Front Microbiol. 2021;12:731008.

51. Stowell SR, et al. Microbial glycan microarrays define key features of host-microbial interactions. Nat Chem Biol. 2014;10(6):470–476.

52. Pinho SS, et al. Immune regulatory networks coordinated by glycans and glycan-binding proteins in autoimmunity and infection. Cell Mol Immunol. 2023;20(10):1101–1113.

53. Saini P, Adeniji OS, Abdel-Mohsen M. Inhibitory Siglec-sialic acid interactions in balancing immunological activation and tolerance during viral infections. eBioMedicine. 2022;86:104354.

54. Kannagi R, et al. Carbohydrate-mediated cell adhesion in cancer metastasis and angiogenesis. Cancer Sci. 2004;95(5):377–384.

55. Haab BB, et al. Definitive Characterization of CA 19-9 in Resectable Pancreatic Cancer Using a Reference Set of Serum and Plasma Specimens. PLoS ONE. 2015;10(10):e0139049.

56. Barnett D, Hall J, Haab B. Automated Identification and Quantification of Signals in Multichannel Immunofluorescence Images The SignalFinder-IF Platform. Am J Pathology. 2019;189(7):1402–1412.

57. Jiang Y, et al. Wilcoxon Rank-Based Tests for Clustered Data with R Package clusrank. J Stat Softw. 2020;96(6). 10.18637/jss.v096.i06.

58. Bates D, et al. Fitting Linear Mixed-Effects Models Using lme4. J Stat Softw. 2015;67(1). 10.18637/jss.v067.i01.

