## Supplementary Text for "Multiplexed Glycan Immunofluorescence Identification of Pancreatic Cancer Cell Subpopulations in Both Tumor and Blood Samples"

Supplementary Information

### Multiplexed Glycan Immunofluorescence Enables the Identification of Divergent Types of PDAC Cell Subpopulations and Tumors

**Braelyn Binkowski*[1], Zachary Klamer*[1], ChongFeng Gao[1], Ben Staal[1], Anna Repesh[1], Hoang-Le Tran[1], David M. Brass[1], Pamela Bartlett[2], Steven Gallinger[3], Maria Blomqvist[4,5], J. Bradley Morrow[2], Peter Allen[6], Chanjuan Shi[6], Aatur Singhi[7], Randall Brand[7], Ying Huang[8], Galen Hostetter[1], and Brian B. Haab[1]**

[1]Van Andel Institute, Grand Rapids, Michigan, USA.
[2]Trinity Health Grand Rapids, Michigan, USA.
[3]University Health Network, Toronto, ON, Canada
[4]Department of Laboratory Medicine, Institute of Biomedicine, University of Gothenburg, Gothenburg, Sweden.
[5]Department of Clinical Chemistry, Sahlgrenska University Hospital, Gothenburg, Sweden
[6]Duke University School of Medicine, Durham, NC, USA.
[7]University of Pittsburgh Medical Center, Pittsburgh, PA, USA.
[8]Fred Hutchinson Cancer Research Center, Seattle, WA, USA.

**Supplementary Tables (in separate Excel file)**

1. Antibody and lectin information
2. Detailed sample information
3. Training set ROI number
4. Test set ROI numbers
5. Complete test set ROI data
6. Test set cancer associations of individual glycan types
7. Results from test set combined classifications
8. Data and thresholds for the tissue-matched blood samples
9. Serum sample information

**Supplementary Methods**

1. Cell culture methods and authentication.

**Supplementary Figures**

1. Representative images of histology annotation types
2. Recursive partitioning to define thresholds and cancer associated signatures
3. Glycan signatures of the cells in association with histology
4. Representative images of cellular glycan signature classifications and ROI percentages in non-cancer and cancer ROIs
5. Determination of tumor within cluster purity and between cluster variance
6. Validation of in-vitro detection of secretions from cell lines

#### Supplementary Methods

##### Cell culture methods and authentication

The PaTu-8988S and PaTu8988T cell lines were obtained from Creative Bioarray (Shirley, NY), and Colo357, L3.3, and L3.6PL lines were kindly provided by Dr. Isaiah J. Fidler (University of Texas, MD Anderson Cancer Center). The remaining cell lines were obtained from ATCC (Manassas, VA). All cell lines were cultured in RPMI-1640 supplemented with 5% fetal bovine serum, 2 mM L-glutamine, and 100 IU/mL penicillin/streptomycin. The cells were grown at 37 °C in a humidified atmosphere supplemented with 5% (v/v) CO2. All cell lines were within 10 passages of collection and use in the described experiments. For preparing chamber slides, cells were seeded onto glass microscope slides that were partitioned into 12 chambers (Ibidi, Lochhamer Schlag, Germany). The cells were seeded at $1 \times 10^4$ cells per chamber and cultured for 3 days using routine culture conditions. The cells were fixed in 10% neutral buffered formalin (Leica, Richmond, IL) for 20 minutes and kept in 1×PBS before proceeding to the multicomplexes immunofluorescence.

Cell line authenticity was confirmed by comparing their RNASeq data to that of published authenticated cell lines, the ATCC Cell Line Land (ACLL), and/or STR profiling (ATCC). Cross contamination between cell lines was excluded by Infinium QC Array (iLLumia, New York, NY) (Genomics core, Van Andel Institute) according to instruction from the provider. In brief, DNA samples of 200 ng were amplified overnight, followed by controlled enzymatic fragmentation. After alcohol precipitation and DNA resuspension, the BeadChip is prepared for hybridization in the capillary flow-through chamber. Samples are applied to prepared BeadChips and incubated overnight. During this overnight hybridization, the DNA samples anneal to locus specific 50-mers covalently linked to up to millions of bead types. One bead type corresponds to each allele per SNP locus. Allelic specificity is conferred by enzymatic base extension followed by fluorescent staining. The iScan System detects the fluorescence intensities of the beads and Illumina software automatically performs analysis and genotype calling. All cell lines were mycoplasma free, as tested by DAPI staining, and proved by RNASeq data that is free of RNA sequence of mycoplasma.

#### Supplementary Figures

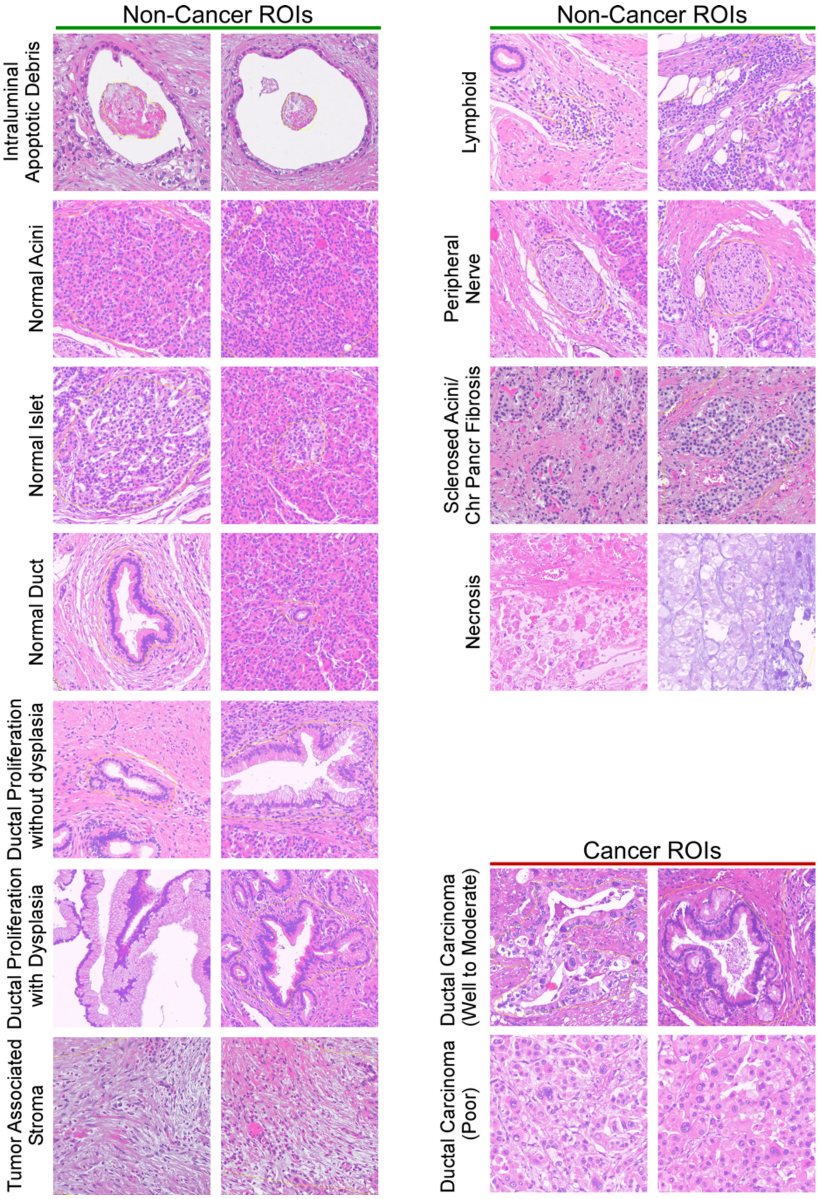


**Figure S1**. Representative images of histology annotation types.

**A**

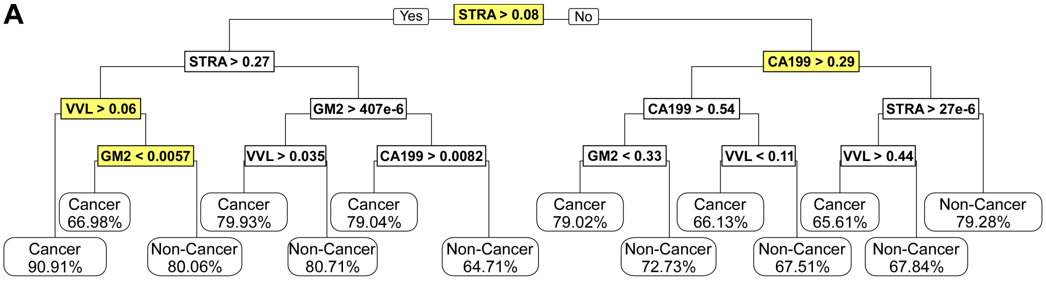


| Importance | STRA | VVL | CA19-9 | GM2 | TRA | GSL-II |
|---|---|---|---|---|---|---|
| Score | 17609.5531 | 5578.0597 | 5398.4080 | 1703.6617 | 1257.7819 | 900.1479 |

**B**

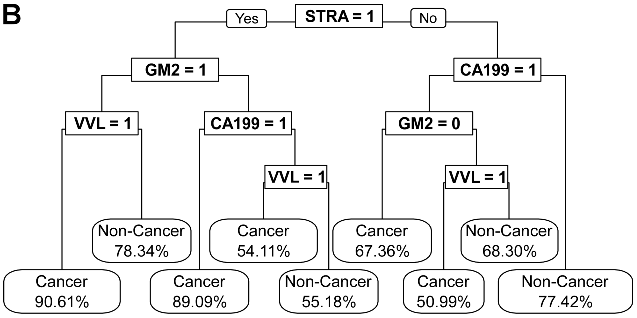


| | Cancer | | Non-Cancer | |
|---|---|---|---|---|
| Glycan Signature | Cell Count | Percent | Cell Count | Percent |
| 0000 | 56360 | 44.07 | 229653 | 63.83 |
| 0001 | 7626 | 5.96 | 53020 | 14.74 |
| 0010 | 1647 | 1.29 | 2348 | 0.65 |
| 0011 | 32 | 0.25 | 1213 | 0.34 |
| 0100 | 4507 | 3.52 | 1189 | 0.33 |
| 0101 | 473 | 0.37 | 681 | 0.19 |
| 0110 | 947 | 0.74 | 158 | 0.04 |
| 0111 | 103 | 0.08 | 267 | 0.07 |
| 1000 | 12618 | 9.87 | 45217 | 12.57 |
| 1001 | 2432 | 1.90 | 9685 | 2.69 |
| 1010 | 9724 | 7.60 | 9879 | 2.75 |
| 1011 | 20914 | 16.35 | 2378 | 0.66 |
| 1100 | 3531 | 2.76 | 3036 | 0.84 |
| 1101 | 561 | 0.44 | 498 | 0.14 |
| 1110 | 4388 | 3.43 | 475 | 0.13 |
| 1111 | 1735 | 1.36 | 84 | 0.02 |

**Figure S2**. Recursive partitioning to define thresholds and cancer associated signatures. A) A classification tree trained on fractional area positive for each glycan for each cell to obtain cutoffs for each glycan. Highlighted nodes denote the cutoffs used for dichotomization. The glycan importance score was calculated from the classification tree. B) The classification tree was trained on dichotomized glycan fractional area per cell to obtain final cancer associated signatures. The table gives the cell counts associated with each signature in Cancer and Non-Cancer ROIs, with percent of cells having the signature.

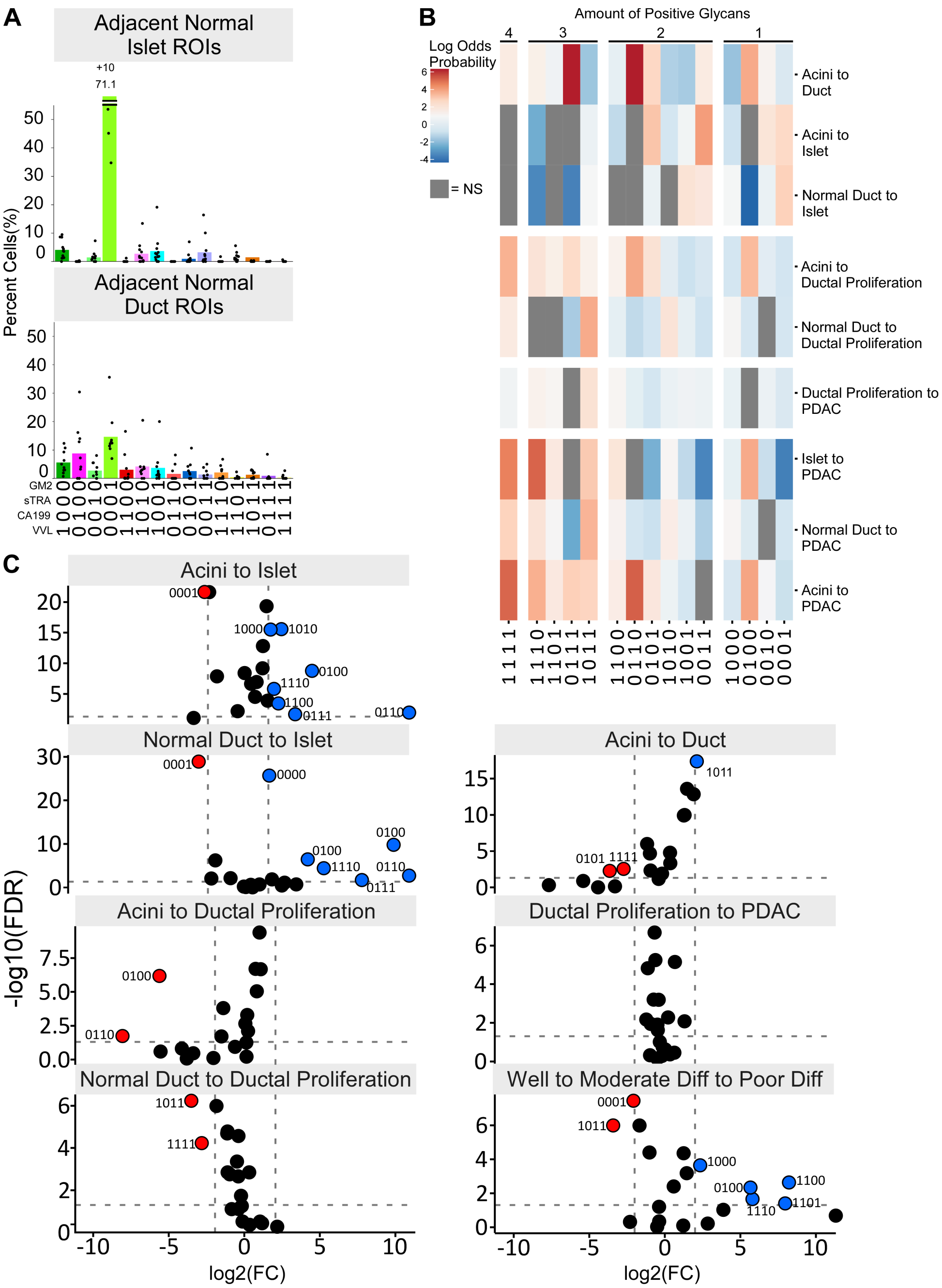



**Figure S3**. Glycan signatures of the cells in association with histology. A) Glycan signature distributions for adjacent normal islets and ducts. B) Volcano plots comparing different cell types. C) Cell probability of belonging to the second cell type compared to the first.

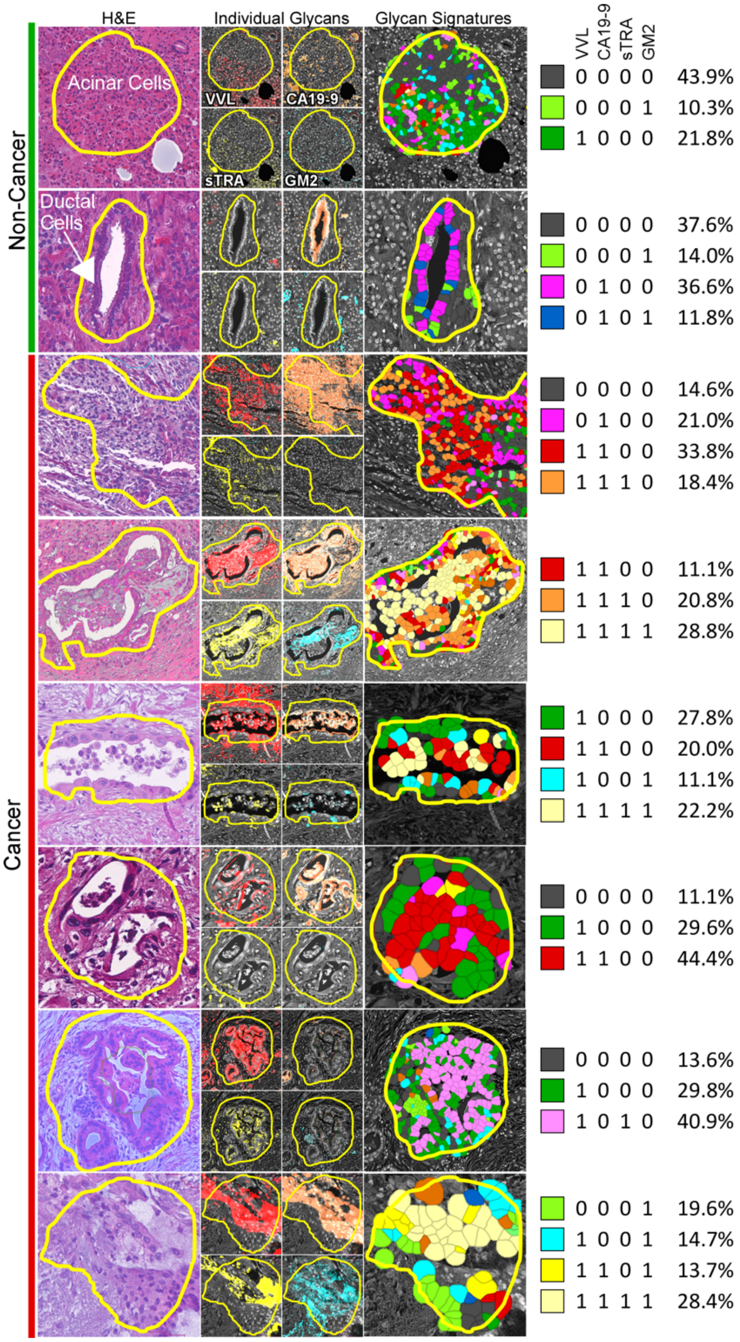


**Figure S4.** Representative images of cellular glycan signature classifications and ROI percentages in non-cancer and cancer ROIs.

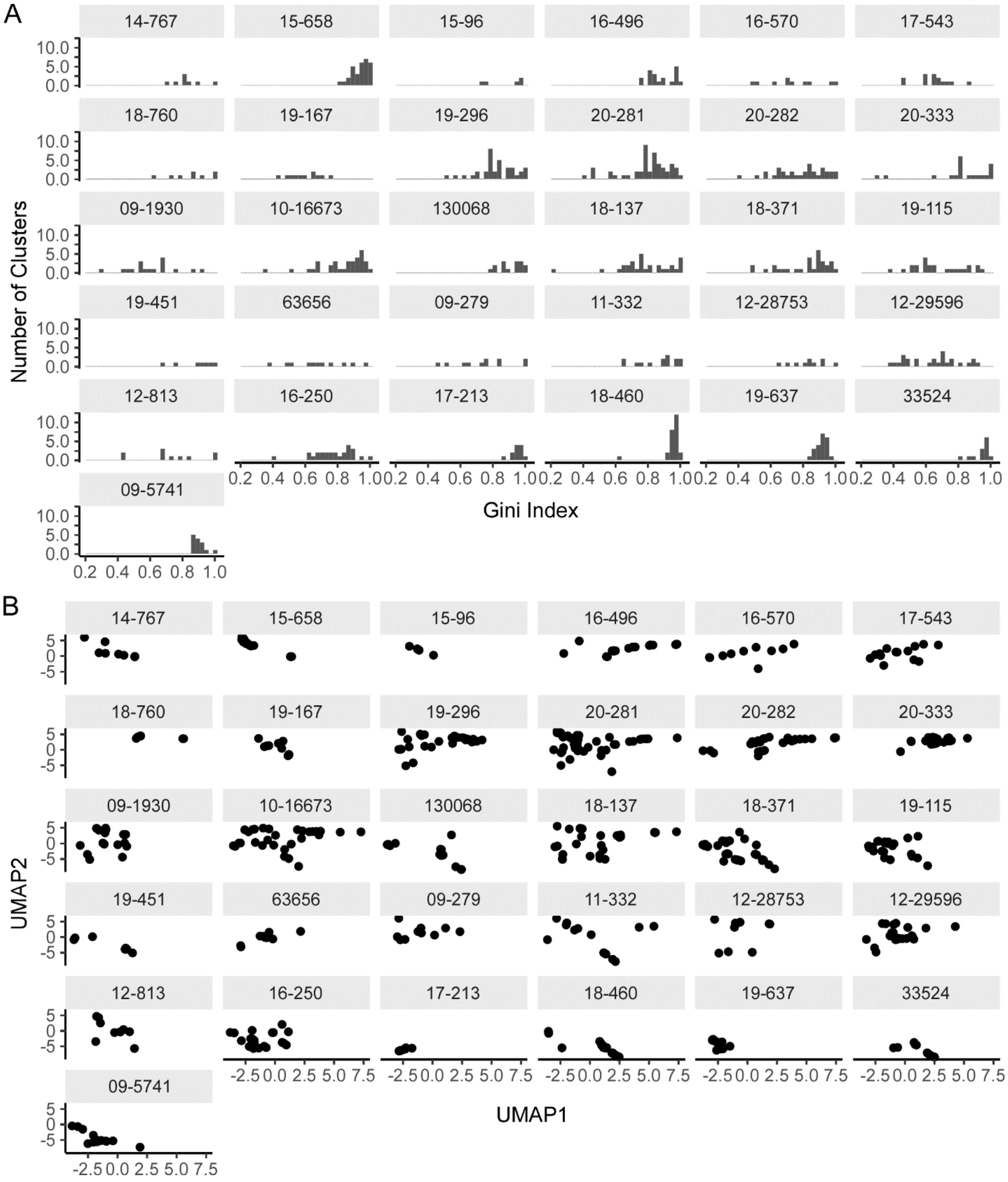


**Figure S5**. Determination of tumor within cluster purity and between cluster variance. A) Per-tumor histograms of cluster Gini index. B) UMAP diagram demonstrating the variation between clusters within tumors. Each graph is data from a single tumor, and each data point in each graph represents a cell cluster. The UMAP diagram was calculated by the similarities between clusters in their percentages of cells belonging to each cancer glycan signature. The average distance between clusters in the UMAP diagram is the average centroid difference between clusters.

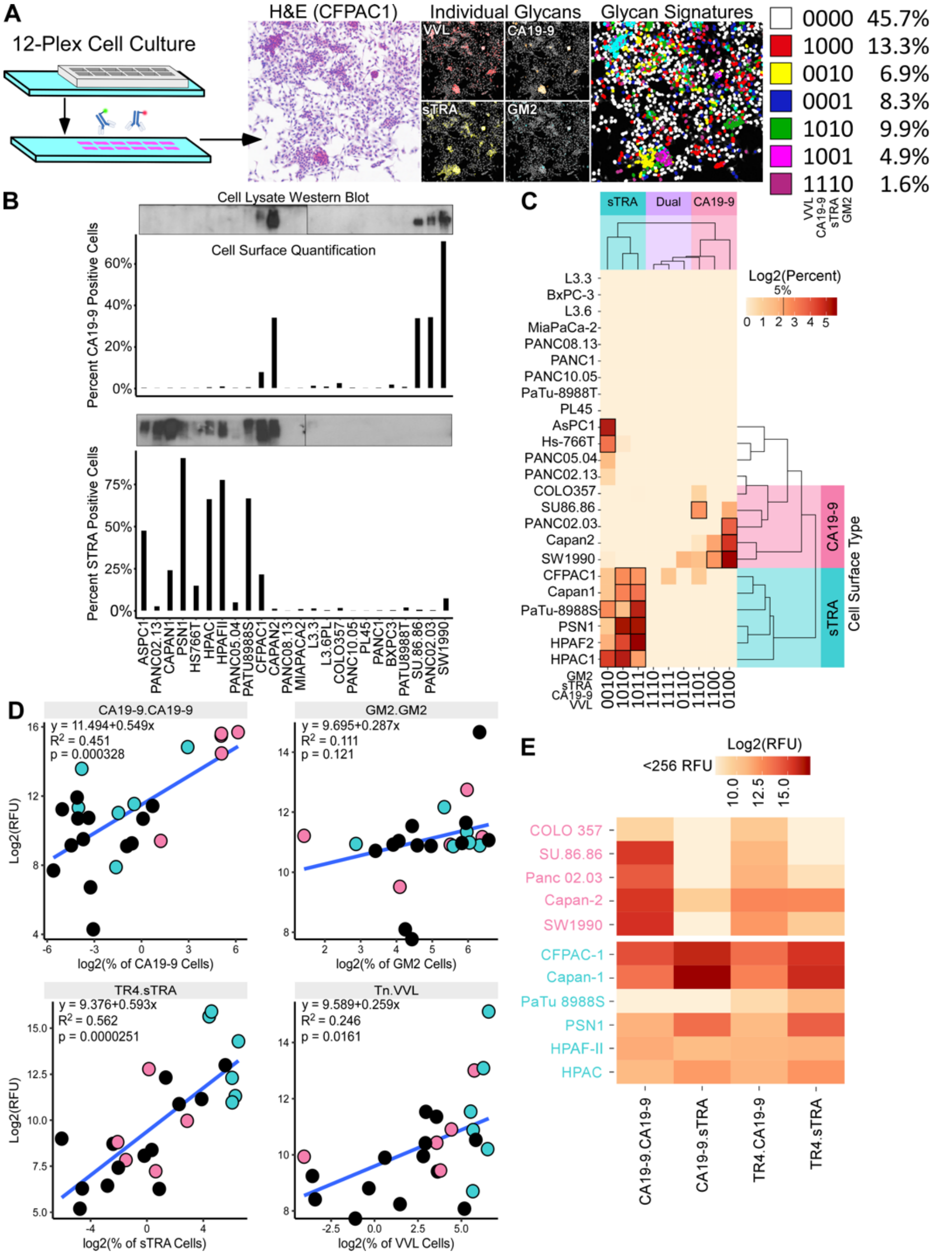


**Figure S6.** Validation of *in-vitro* detection of secretions from cell lines. A) Methodology and example data, using equivalent methods as described in the main text for the primary tumor specimens. B) Comparison of Western blot data with the quantification of cell counts using immunofluorescence. The column graphs give the percentage of cells in each cell line that were positive for CA19-9 (top) or sTRA (bottom). C) Variation in glycan signatures between cell lines. Each value in the matrix is the percentage of cells in each cell line with each of the indicated glycan signatures. D) Correlation between cell surface and secretion. The y-axis gives relative fluorescence of the indicated media assay, and the x-axis gives the percentage of cells in each cell line with the indicated glycans. Each datapoint is a cell line, color coded by the cell surface type given in panel C. E) Cell line conditioned media assay response.
